# Ultrastructural analysis of neuroimplant-parenchyma interfaces uncover remarkable neuroregeneration along-with barriers that limit the implant electrophysiological functions

**DOI:** 10.1101/2021.10.03.461535

**Authors:** Aviv Sharon, Nava Shmoel, Hadas Erez, Maciej M. Jankowski, Yael Friedmann, Micha E. Spira

## Abstract

Despite increasing use of in-vivo multielectrode array (MEA) implants for basic research and medical applications, the critical structural interfaces formed between the implants and the brain parenchyma, remain elusive. Prevailing view assumes that formation of multicellular inflammatory encapsulating-scar around the implants (the foreign body response) degrades the implant electrophysiological functions. Using gold mushroom shaped microelectrodes (gMμEs) based perforated polyimide MEA platforms (PPMPs) that in contrast to standard probes can be thin sectioned along with the interfacing parenchyma; we examined here for the first time the interfaces formed between brains parenchyma and implanted 3D vertical microelectrode platforms at the ultrastructural level. Our study demonstrates remarkable regenerative processes including neuritogenesis, axon myelination, synapse formation and capillaries regrowth in contact and around the implant. In parallel, we document that individual microglia adhere tightly and engulf the gMμEs. Modeling of the formed microglia-electrode junctions suggest that this configuration suffice to account for the low and deteriorating recording qualities of in vivo MEA implants. These observations help define the anticipated hurdles to adapting the advantageous 3D in-vitro vertical-electrode technologies to in-vivo settings, and suggest that improving the recording qualities and durability of planar or 3D in-vivo electrode implants will require developing approaches to eliminate the insulating microglia junctions.

## Introduction

Basic and clinically oriented brain research and their applications rely on the use of sophisticated neuro-implants for long-term, simultaneous, multisite extracellular recordings of field potentials (FP) generated by neurons in freely behaving subjects. Despite significant technological progress, contemporary in-vivo multielectrode array (MEA) technologies suffer from inherent limitations that include: (a) a low signal-to-noise ratio (S/N), (b) low source resolution and (c) deterioration of the recording yield and FP amplitudes within days to weeks of implantation (Jackson and Fetz, 2007;Perge et al., 2013;Voigts et al., 2013;Harris et al., 2016;Lee et al., 2018;Lee et al., 2021). In addition, current in-vivo brain implants are “blind” to sub-threshold synaptic potentials generated by individual neurons. This implies that critical elements of the brains signaling repertoire and computational components are ignored. The prevailing view relates these limitations to: (a) the gradual increase in the thickness of the inflammatory glia scar that displaces neurons from the implant surfaces (Edell et al., 1992; Biran et al., 2005;Polikov et al., 2005;Malaga et al., 2016;Salatino et al., 2017a;Michelson et al., 2018), (b) the glial scar encapsulating the implant (Szarowski et al., 2003;Johnson et al., 2005;Polikov et al., 2005;Otto et al., 2006;Williams et al., 2007;Prasad and Sanchez, 2012) and a biofouling layer assembled on the electrode surfaces insulate the electrodes from the current sources by their relatively high resistivity compared to the intact brain tissue (Sommakia et al., 2009;Sommakia et al., 2014;Malaga et al., 2016), (c) pro-inflammatory cytokines released from the glia and injured neurons lead to demyelination of the axons and thereby disrupt action potential propagation (Winslow et al., 2010;Winslow and Tresco, 2010), (d) released cytokines reduce the excitability and synaptic connectivity of neurons in the implant’s vicinity (Vezzani and Viviani, 2015;Salatino et al., 2017b;Hermann and Capadona, 2018;Salatino et al., 2019;Thompson et al., 2020), (e) damage to blood capillaries by the implant leads to infiltration of neurotoxic factors and myeloid cells (Saxena et al., 2013) and reduces the blood supply to individual cells. Although objective experimental attempts to relate the thickness of the inflammatory foreign body response (FBR) to deterioration in recording qualities have failed, this concept has continued to dominate the field and still shapes extensive research efforts to mitigate or overcome this deterioration. Whereas ever-improving spike-detecting, spike-sorting and signal averaging techniques make it possible to extract significant information from monitoring extracellular FP (Quiroga et al., 2004;Einevoll et al., 2012;Carlson and Carin, 2019), the limited recording qualities of current multielectrode array-implants (MEA implants) and their deterioration in time considerably hinder the research progress.

The realization that the use of substrate integrated planar MEA technologies for extracellular recordings (Figure 1) inherently limits the qualities of in-vitro and in-vivo systems has prompted the development of new 3D in-vitro technologies to enable parallel, multisite intracellular recordings and stimulation from many individual cultured cells (neurons, cardiomyocytes and striated muscles). In principle, this family of in-vitro MEA technologies utilizes different forms of 3D vertical nano-structures (nano-pillars) that pierce the plasma membrane of cultured cells (by electroporation or spontaneously) in a way similar to classical sharp electrodes (Figure 1 and Tian et al., 2010;Angle and Schaefer, 2012;Duan et al., 2012;Gao et al., 2012;Robinson et al., 2012;Xie et al., 2012;Angle et al., 2014;Lin and Cui, 2014;Lin et al., 2014;Qing et al., 2014;Abbott et al., 2017;Dipalo et al., 2017;Liu et al., 2017;Abbott et al., 2018;Abbott et al., 2019;Mateus et al., 2019;Li et al., 2020;Teixeira et al., 2020;Yoo et al., 2020;Xu et al., 2021;Zhang et al., 2021).

**Figure 1.**
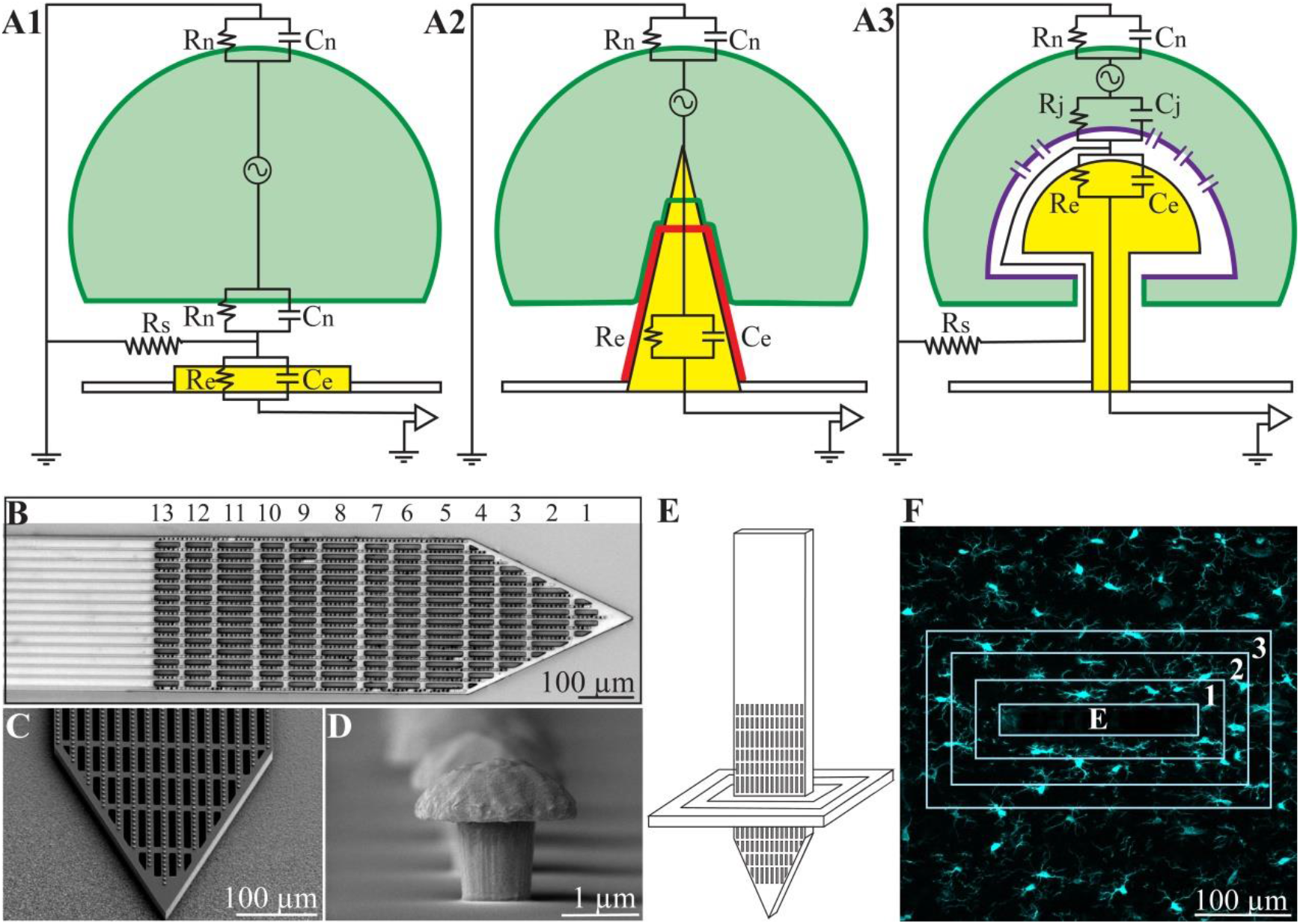
(A) Passive analog electrical circuit models depicting the structural interfaces formed between cells and recording microelectrodes under in-vitro conditions. (A1) substrate integrated planar electrode for extracellular field potential recordings, (A2) a vertical nano-pillar electrode that pierces the plasma membrane for intracellular recordings, and (A3), extracellular gold mushroom shaped vertical microelectrode (gMμE) for IN-CELL recordings. The three configurations differ mainly in terms of the nature and dimensions of the cleft formed between the cultured cells (neurons, cardiomyocytes or striated muscle fibers) and the recording electrode. In (A1), the extracellular field potential generated by propagating action potentials is largely attenuated across the high resistance non-junctional membrane (Rn) and the low seal resistance (Rs). In (A2), the vertical nanoelectrode pierces the cell’s plasma membrane, gaining direct access to the cytosol (Rn is reduced to zero). A very high seal (∼GΩ) resistance (not drawn) formed between the vertical nanoelectrode’s (yellow electrode with a red insulating layer) surface and the plasma membrane (green). In (A3), the cell engulfs a mushroom shaped vertical electrode (yellow) to form relatively high Rs by the narrow cleft. Along with reduced junctional membrane resistance (Rj -purple) the configuration makes it possible to record attenuated intracellular potentials. (B) Low magnification image of the polyimide based perforated MEA platform (PPMP), the proximal solid part and distal perforated part are shown. For orientation, the rows of perforations are numbered. (C) SEM enlargement of a PPMP segment, showing the perforations of the polyimide platform and dense rows of gMμEs along the PI “ribs”. (D) A SEM image of a gMμE. (E) Schematic illustration of an implanted PPMP and the orientation of thick horizontal tissue slices. (F) The integrated immuno-fluorescent intensity within the electrode (central rectangle-E) and within 25 μm wide centripetal shells around it were measured and processed to establish the normalized fluorescent intensity level (NFI) or the number of a given cell type at a given distance and time around the implant.

At the same time, a number of laboratories have developed the “IN-CELL” recording and stimulation configuration, in which micrometer-sized, extracellular gold mushroom-shaped microelectrodes (gMμEs) record attenuated synaptic and action potentials (Figure 1 and Spira et al., 2007;Hai et al., 2010b;a;Fendyur and Spira, 2012;Spira and Hai, 2013;Rabieh et al., 2016;Shmoel et al., 2016;Weidlich et al., 2017;McGuire et al., 2018;Spira et al., 2018;Mateus et al., 2019;Spira et al., 2019;Jones et al., 2020;Teixeira et al., 2020). Ultrastructural imaging complemented by electrophysiology and model system analysis of the cultured-neurons/gMμEs configuration have revealed that the biophysical principles of “IN-CELL” recordings are identical to those of the perforated patch electrode configuration (Horn and Marty, 1988;Akaike and Harata, 1994).

Successful adaptation of the vertical nano-pillar and gMμEs MEA approaches to in-vivo brain research could effectively address the limitations of the currently used planar MEA technologies (low S/N, poor source resolution and deterioration), and importantly would make it possible to record the entire signaling repertoire from many individual neurons. It is thus expected that such adaptation will significantly improve the likelihood of understanding the codes of brain-circuit computations.

Ultrastructural examinations of the interfaces formed between cultured neurons and gMμEs or vertical nano-pillar based MEAs have played key roles in revealing that cultured neurons and other cell types tightly engulf vertical structures by evolutionarily conserved cell biological mechanisms (Hai et al., 2010b;Santoro et al., 2014;Santoro et al., 2017b;McGuire et al., 2018). And, that the narrow cleft formed between the engulfing plasma membrane and the gMμEs form a high seal resistance (Rs). This, together with the increased conductance of the cell’s membrane that faces the gMμEs (the junctional membrane – Rj, Figure 1), make it possible to record attenuated action potentials and subthreshold synaptic potentials with features and biophysics similar to perforated patch recordings (Horn and Marty, 1988;Akaike and Harata, 1994;Spira et al., 2007;Hai et al., 2009a;Hai et al., 2009b;Fendyur et al., 2011;Santoro et al., 2013;Santoro et al., 2014;Santoro et al., 2017a;Santoro et al., 2017b).

In contrast to meticulous ultrastructural studies of the interfaces formed between cultured cells and different types of vertical nanoelectrodes, structural studies of the interfaces formed between implanted neuroprobes and in-vivo brain parenchyma were of very low resolution. Besides the inherent low spatial resolution of the immunohistological methods used, in the vast majority of light and electron microscope studies, the implants were pulled out (extracted) from the brain tissue prior to thin sectioning for histological examination. This unavoidably damages the parenchyma/implant interfaces, making it impossible to examine and understand the structural relationships between the abiotic implant and the tissue (for example Schultz and Willey, 1976;Moss et al., 2004;Grand et al., 2010;Marton et al., 2020).

Using gMμEs based perforated polyimide MEA platforms that can be thin sectioned along with the interfacing parenchyma; we examined here for the first time the interfaces formed between brains parenchyma and implanted 3D vertical microelectrode (gMμEs) platforms at the ultrastructural level. Our study demonstrates remarkable structural parenchyma regenerative processes including neuritogenesis, axon myelination and synapse formation in contact and around the implant. In parallel, we documented that individual microglia adhere tightly and engulf the gMμE electrodes. The extracellular cleft formed between the implant and the adhering microglia in parallel to the microglia’s input resistance suggest that high resistance barriers are formed in contact with the electrodes. We posit that these microglia-electrode-junctions, rather than the thick multicellular inflammatory encapsulation that is thought to displace neuronal cell bodies and induce axon demyelination or structural synapse degeneration, are the underlying mechanisms governing the deterioration of the electrical coupling between neurons and the in-vivo implanted electrodes. In addition, our ultrastructural observations objectively highlight the expected hurdles to applying arrays of vertical nano-pillars in general and gMμEs in particular to record intracellular potentials from cortical neurons in freely behaving rats. Approaches to mitigate or selectively eliminate the adhering microglia are thus needed to advance the application of 3D microelectrode arrays for intracellular recording of the entire signaling repertoire of the in-vivo brain.

## Materials and Methods

### Animals

All the procedures in the present study were approved by the Committee for Animal Experimentation at the Institute of Life Sciences of the Hebrew University of Jerusalem. This study was conducted using female Sprague Dawley rats (215-340 g).

### Neuro-implants

To address the technical features required to prepare thin sections of implanted gMμE-platforms along with the parenchyma around it, we fabricated nonfunctional implants constructed of a Perforated Polyimide (PI)-based MEA Platform (PPMP) that carries a dense array of gold mushroom shaped microelectrodes (gMμE, Figure 1). The 1.7 mm long, 280 μm wide and 16 μm thick nonfunctional gMμE-PPMPs were divided to a 0.9 mm long solid proximal part and a 0.8 mm perforated distal part (Figure 1). The perforated segment tapered to form a sharp tip. The width of all the rectangular perforations was 7-8 μm and the lengths of the different perforations were 65, 47 and 44 μm (Figure 1). gMμEs were electroplated at a pitch of 8 μm in rows along the15 gold conducting lines that run along the platform (Figure 1 and Supplementary Figure S1).

### Implant fabrication

The gMμE-PPMPs were constructed using standard photolithography fabrication methods as follows (Supplementary Figure S1). First, an aluminum releasing layer was sputtered on a 3-inch silicon wafer (University Wafer, USA), followed by a spin-coated 15 μm thick polyimide layer (PI 2610, HD Microsystems, Germany) that served as an insulating layer and the main mechanical backbone of the platform. A triple metal layer of Cr/Au/Cr (20/120/20 nm) was then patterned and e-beam evaporated as interconnects, pads and scribe-lines. Next, a second 1 μm thick insulating layer of polyimide was spin-coated, followed by the deposition of a 1 μm SiO_2_ with Plasma Enhanced Chemical Vapor Deposition (PECVD). A 1.5 μm photoresist layer was then patterned. Dry etching by RIE was used to define 1.5 μm molds for electroplating the gMμEs and pads through the SiO_2_ and the one micrometer thick insulating PI layer. After removal of the top Cr layer by wet etch, gMμEs with cap diameters of ∼2 μm were electrodeposited at a pitch of 8μm along the tip and a perforated section of the platform (Figure 1 and Supplementary Figure S1). An additional 300 nm of SiO_2_ was deposited with PECVD and a photoresist layer were used to define the perforated pattern of the platform. The photoresist and SiO2 layers were then removed and the platforms were then released from the wafer by anodic metal dissolution and thoroughly rinsed in distilled deionized water.

### Platform implantations

A 1-1.5 cm longitudinal cut of the skin on the head was made and the anterior, dorsal surfaces of the skull were exposed. Two craniotomies, one in the left and the other in the right frontal bones, were performed at the desired reference points (coordinates: AP: +3.5 mm; ML: +2.5 mm from the Bregma) and the dura was gently resected (0.3-0.5 mm long incision). The 1.7 mm long platforms held by forceps mounted on a micromanipulator were slowly inserted into the motor cortex to a depth of 1.8 mm. The electrodes were gently released from the holder and the craniotomy was sealed with melted bone wax (W810, Ethicon, Belgium). The wound was treated in situ with antibiotic ointment (Synthomycine, chloramphenicol 5%) and sutured with nylon sutures. Then the rats received an intraperitoneal injection of the antibiotic Enrofloxacin 50 mg/ml (5% W/V) at a dose of 15 mg/kg diluted with saline to 1 ml (Baytril, Bayer Animal Health GmbH, Leverkusen, Germany). In line with standard protocols to prevent postoperative pain, the rats received for three consecutive days after gMμE-PPMP implantation non-steroidal anti-inflammatory/analgesic drugs. A subcutaneous injection of Carprofen 50 mg/ml (5% W/V) in a dose of about 12 mg/kg (Norocarp, Norbrook Laboratories Limited, Newry, Co. Down, Northern Ireland) during surgery. To further reduce the stress and pain caused by injections and prevent mechanical stress to the skin around the implantation site, the rats were fed on days 2 and 3 post PPMP implantation by Meloxicam (Rheumocam, oral suspension 1.5mg/ml, Chanelle pharma) dissolved in palatable Jelly. To that end, Meloxicam dissolved in agar (Meloxicam-jelly) prepared in a small Petri dish (diameter of 35 mm) was placed in the rat cages. The Petri dishes were removed at the end of days 2 and 3. Visual checks confirmed that the rats consumed the entire volume of the Meloxicam-jelly. After surgery, the animals were housed individually to prevent them from chewing the implants.

### Tissue processing for immunohistology and transmission electron microscopy

For brain tissue fixation, individual rats were deeply anesthetized with isoflurane (Piramal, United States) followed by an IP overdose injection of pentobarbital (4.5ml per 250g rat, CTS Group, Israel). When breathing had stopped, the rats were transcardially perfused with phosphate buffer saline (PBS). This was followed by a 4% paraformaldehyde in PBS (PFA, Sigma-Aldrich) perfusion for immunohistology and 1-2.5% glutaraldehyde/2% paraformaldehyde (Agar Scientific) for transmission electron microscopy (TEM). In both cases, the perfusion rate was 10 ml/min and lasted for 40 min. Next, the skulls were removed and the implanted brains were post-fixed at 4°C for an additional 12-24 hrs. in either PFA (for immunohistology), or glutaraldehyde/paraformaldehyde (for TEM). Thereafter, the fixed and exposed brains destined for immunohistology were washed in PBS and incubated for 1-3 days in a 30% sucrose solution in PBS at 4°C.

To prepare the brain tissue for cryosectioning (immunohistology), cubic shaped portions of tissue (approximately 1×1×1cm) with the PPMP in their center were isolated. The isolated piece was placed in a freezing medium (Tissue-Plus O.C.T. Compound, Scigen) and frozen at -80°C. The frozen tissues along with the implanted platform were then horizontally sectioned into 40 μm thick slices using a Leica CM1850 Cryostat. Individual slices were collected and placed in 24 well plates containing PBS. The tissue slices were then incubated in blocking solution (1xPBS, 1% heat-inactivated horse serum (Biological Industries), 0.1% Triton X-100 (Sigma Aldrich)) for 1 hour at room temperature (RT) under gentle shaking. Next, the slices were incubated with a diluted primary antibody for 3 hours at room temperature (RT) and washed 3 times with the blocking solution. This was followed by 1-hour incubation at RT with the diluted secondary antibody after which the slices were washed with the blocking solution 3 times and stained with the nuclear marker DAPI (Sigma–Aldrich, 1 mg/ml 1:1000) for 15 min at RT. After washing with the blocking solution and PBS, the slices were mounted on Superfrost Plus Slides (Thermo Fisher Scientific) and sealed by a Vectashield (VE-H-1000 -Vector Labs) mounting medium.

### Electron microscopy

For TEM imaging, glutaraldehyde/paraformaldehyde fixed tissue along with the PI, MEA platform implants were washed with PBS and sliced by a Leica VT1000S Vibrotome using a ceramic blade (Campden Instruments Ltd.) into 200 μm thick horizontal sections. The slices were deposited in 24 well plates with PBS.

After 8 washes with 0.1 M cacodylate buffer at pH 7.4 (SigmaAldrich) the tissue was post fixed by 1% osmium tetroxide (Electron Microscopy Sciences) and 0.6% K3Fe(CN)_6_ in a 0.1 M cacodylate buffer for 1 hr. at room temperature. The slices were then washed again in a 0.1 M cacodylate buffer and dehydrated by a series of increasing concentrations of ethanol solutions of 10%, 25%, 50%, 75%, 90%, 96%, 100%, 100%. Finally, the slices were embedded in Agar 100 (Agar Scientific). The embedded preparation was then thin-sectioned and observed using a TEM Tecnai 12 microscope at 100 kV. The shown TEM images were taken from sections prepared across the perforated part of the implant (rows 2-6 as marked in Figure 1). Efforts were made to orientate the thin sections perpendicular to the long axis of the polyimide platform and along the gMμEs. As the diameter of the stalks and caps of the gMμEs are in the range of 1-3 μm, a slight deviation from a perfect sectioning angle, resulted in imperfect sections that did not pass through the entire length of the gMμE cap and stalk. Thus, in some cases, the sections went through the entire length of the mushrooms cap, stalks and the contact between the stalk and the conducting line (for example Figures 6 and 7A). In others, the thin sections were slightly tilted in respect to the long axis of the gMμE stalks. In these cases the stalk of the mushroom appears to taper toward the polyimide platform. The observed TEM images and conclusions represent transmission electron microscope imaging of over >250 thin sections prepared from 25 different PPMPs implants.

### Immunolabeling, confocal imaging, image processing, analysis and statistics

Immunolabeling, imaging, image processing and analysis were conducted as detailed in previous studies from our laboratory (Huang et al., 2020;Sharon et al., 2021). Briefly, neurons were concomitantly labelled with two antibodies: one for neurite labelling (mouse anti-Neurofilament 160/200 monoclonal antibody (Sigma-Aldrich N2912, 1:10000-1:20000) and the other for neuronal nuclei (mouse anti-NeuN monoclonal antibody (Merck MAB377, 1:200)). Astrocytes were labelled with chicken anti-glial fibrillary acidic protein (GFAP) polyclonal antibodies (Thermo Fisher PA1-10004, 1:500-1000). Microglia were labelled using rabbit anti-Iba-1 monoclonal antibody (Abcam ab178846, 1:1000). For the secondary antibodies we used goat anti-mouse Alexa 488, goat anti-chicken Alexa 647 (Thermo Fisher A-11001 and A21449 respectively, 1:100) and sheep anti-rabbit Cy3 (Sigma–Aldrich C2306, 1:100).

Confocal image stacks of the immunolabeled slices were acquired with an Olympus FLUOVIEW FV3000 confocal scan head coupled to an IX83 inverted microscope, using a 20X air objective (NA= 0.75). Typically, 15-30 confocal slices were acquired, with a vertical spacing of 1μm. Image processing of the immunolabeled sections was conducted using the Fiji distribution of ImageJ (Schindelin et al., 2012;Schneider et al., 2012).

Two methods of analysis and representation of the cell densities in contact and around the PPMPs were used: (1) The densities of the astrocytes and neurons, including their cell bodies and neurites, were analyzed and displayed as the relative fluorescent intensities with respect to the normal background. These are referred to in Figure 2 E2 and E3 as the Normalized Fluorescent Intensity (NFI) values (Huang et al., 2020). (2) The density of the microglia and neuronal cell bodies per 100 μm^2^, at a given shell around the implant, and within the pores were calculated by manual counting (Figure 2 E1 and E4). The counting of these cell bodies was done by merging Iba1 labelled microglia or NeuN labeled neurons with the nuclear marker DAPI.

**Figure 2.**
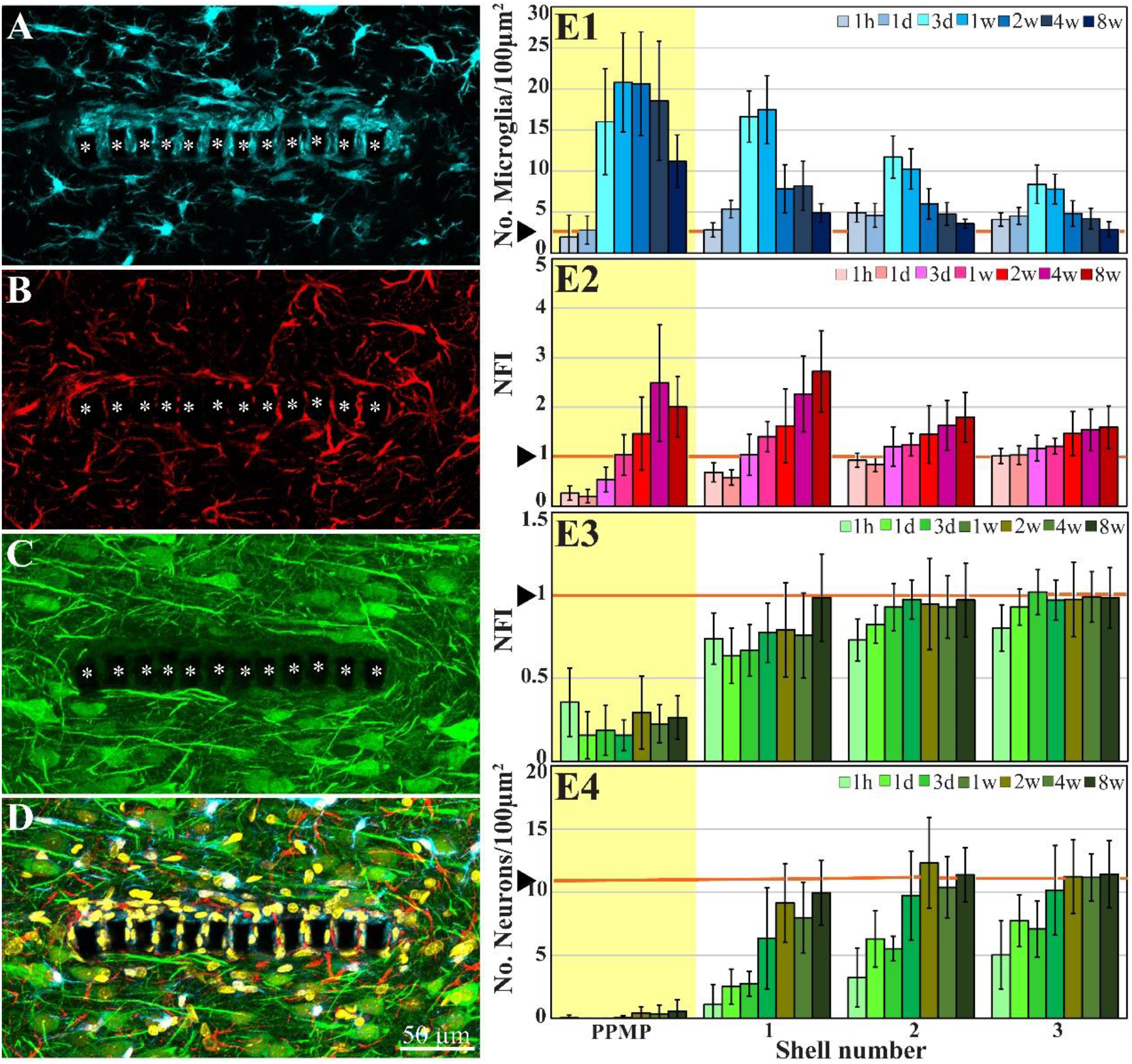
Confocal microscope images showing horizontal-sections of immuno-labeled cortical brain tissue along with cross sections of the perforated segment of an implanted PPMP, two weeks post implantation. Shown are: microglia (cyan) within and around the implant marked by asterisks (A), astrocytes (B, red), neurons and neurites (C, green), and a merged image of (A-C) which also includes the nuclei of the cells labeled in yellow (D). (E) Histograms depicting the average Normalized Fluorescent Intensity (NFI) or number of cells/100μm^2^. Microglia (E1, cyan), astrocytes (E2, red), neurites and cell bodies (E3, green) and neuronal cell bodies (E4, green) within and around the platform’s perforated segments. The time post-platform implantation is coded by the darkening of the column color as indicated by the legend on the right hand side of the histograms. The average NFI values or the cells/100 μm^2^ within the platforms (PPMP) are highlighted in yellow. The distance of the average NFI from the MEA platform is given by shell number. Each shell is 25μm wide (as illustrated in Figure 1F). Vertical lines correspond to one standard deviation. The orange lines indicated by the arrowheads depict the normal NFI values or the number of cell/100μm2 in the control cortices. An enlarged image of D is presented as Supplementary Figure S2.

Average fluorescent values and cell counting characterizing the FBR in space and time were measured and calculated from cortical brain slices prepared from sections from across rows 5-8 within the 280 μm wide, perforated part of the implant (Figures 1B and 2). We used 2-10 hemispheres/experimental points in time (Supplementary Table 1). Each brain hemisphere was used to prepare 1 to 6 tissue slices. Each slice was used to prepare a single maximal projected image generated by 10 consecutive optical sections. For more data on the numerical values and statistical tests, see Supplementary Table 2.

## Results

### Probe design principles

To address the technical features required to prepare thin sections of implanted gMμE platforms along with the parenchyma around it, we fabricated nonfunctional implants constructed of a Perforated Polyimide (PI)-based MEA Platform (PPMP) that carries a dense array of gold mushroom shaped microelectrodes (gMμE-PPMP, Figure 1). PI was selected because it is a biocompatible polymer with a Young’s modulus of 2.5 GP. Importantly, based on studies demonstrating that PI implants can be thin-sectioned for histological examinations (Mercanzini et al., 2007;Mercanzini et al., 2008;Richter et al., 2013;Xie et al., 2014;Boehler et al., 2017;Huang et al., 2020) our laboratory has developed procedures to section implanted gMμE-PPMPs along with the surrounding brain parenchyma for light and transmission electron-microscope (TEM) studies (Huang et al., 2020;Sharon et al., 2021). In the present study, we fabricated 1.7 mm long, 280 μm wide and 16 μm thick nonfunctional gMμE-PPMPs. The proximal 0.9 mm of the implant was constructed of solid PI, and the remaining distal part was perforated (Figure 1). The perforated segment tapered to form a sharp tip. The width of all the rectangular perforations was 7-8 μm and the lengths of the different perforations were 65, 47 and 44 μm (Figure 1). gMμEs were electroplated in rows along the conducing gold lines which run in between the perforations (Supplementary Figure S1). The high density of the gMμEs served to increase the probability of successfully preparing thin sections (80 nm) for TEM imaging through gMμEs and PI along with the interfacing brain parenchyma. The perforated microarchitecture of the platform reduced the projected solid surface area of the perforated part by 35% and allowed cells to extend branches or migrate through the perforations. Each pore in the PI platform approximately doubled the PI surface to which the cells could adhere.

### Ultrastructure of the implant and parenchyma

To examine the interfaces formed between the brain parenchyma and implanted gMμE-PPMPs, cross-sections for transmission electron microscopy of gMμE-PPMPs along with the surrounding tissue were prepared. We selected to examine the ultrastructure of the inflammatory scar at two, four and eight weeks after electrode implantation, since our earlier immunohistological studies showed characteristic alterations in the distribution and densities of the microglia, astrocytes and neurons at these points in time (Figure 2 and Huang et al., 2020;Sharon et al., 2021). For the reader’s convenience, the overall spatiotemporal relationships between microglia, astrocytes, neurons and PPMP implants is briefly presented in Figure 2 using conventional immunohistological imaging.

For the ultrastructural analysis, gMμE-PPMP implanted brains were chemically fixed by standard transcardial perfusion of glutaraldehyde/paraformaldehyde fixative. Since MEA platform implantation unavoidably damages blood capillaries along the insertion path, concern was raised whether the quality of tissue fixation around the implant will suffice to preserve the tissue ultrastructure. In retrospect, based on the preservation qualities of the cell membranes and subcellular organelles including the mitochondria, the smooth and rough endoplasmic reticulum, synaptic vesicles, post-synaptic densities and myelin, we concluded that the perfusion of the fixative was not impaired in the surroundings of the implant. It is important to note, however, that as in other ultrastructural studies of the CNS, the extracellular spaces between the various cell types is reduced by approximately 20% (Korogod et al., 2015;Hrabetova et al., 2018;Soria et al., 2020). Since the volume of the implanted PPMPs is not altered by the fixatives, transcardial fixation led to the generation of mechanical tension around the implant. This often tears the tissue around the implant. Importantly, tissue growing into the PPMP pores and adhering to the platform surfaces remained tightly attached to the platforms, and the break in the tissue took place between cells or even across cells a few micrometers away from the implant surface.

Since the relative positioning of the PPMP and the cells around it are not altered by the classical method of transcardial fixative perfusion, the TEM analysis presented here suffices to provide the essential and missing information on implant brain-tissue interfaces. In addition, because the range of the shrinkage factor is known, the genuine extracellular clefts can be estimated. It is important to note that TEM examination of hundreds of thin sections representing over 25 gMμE-PPMP implants revealed that the gMμEs maintain stable contact with the conducting lines used for their electroplating. That is, the gMμEs are not striped off during the platforms insertion or during the thin sectioning of the tissue along with the implant for TEM imaging.

### Insulation of the PPMP implants by microglia at two weeks post PPMP implantation

Typically, at two weeks post gMμE-PPMPs implantation, tightly adhering dark cytoplasmic microglia processes (dark as compared to other cell profiles in their surroundings) encapsulated individual PI “ribs” (Figures 3 and 4). The cell bodies from which the dark cytoplasm emanated contained characteristic microglia nuclei with clumps of heterochromatin beneath the nuclear envelope and throughout their nucleoplasm (Figure 3 and see Tremblay et al., 2012;Garcia-Cabezas et al., 2016;Savage et al., 2018;Nahirney and Tremblay, 2021). The electron-dense cytoplasm of these microglia was bordered by a clear plasma membrane and contained rough endoplasmic reticulum, mitochondria and dark inclusions which may plausibly be lysosomes and lipofuscin granules (Figures 3 and 4).

**Figure 3.**
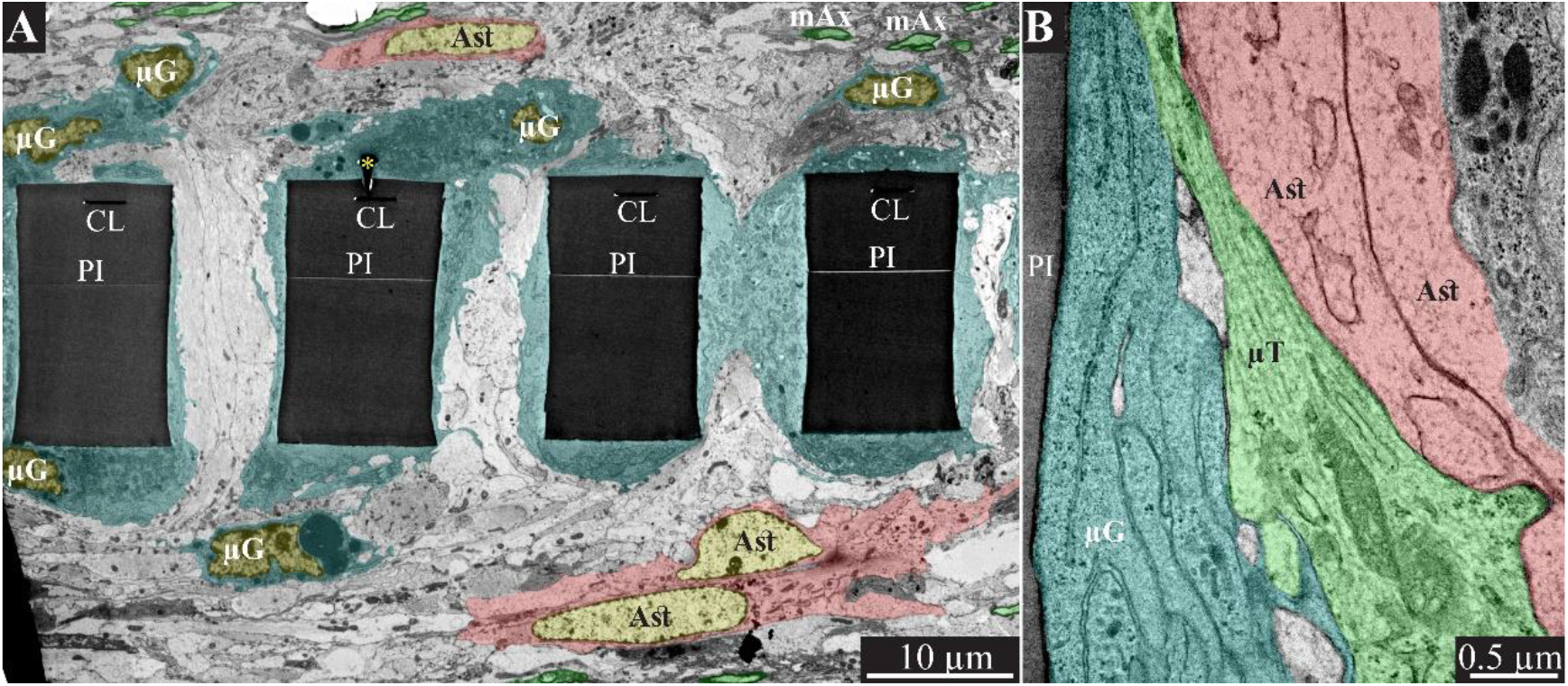
Encapsulation of individual PPMPs “ribs” by adhering microglia. (A) A low magnification transmission electron microscope cross-section of an implanted PPMP along with the parenchyma around it 2 weeks after platform implantation. The PI “ribs” are encompassed by microglia (cytoplasm marked in cyan and typical microglia nuclei μG - yellow) and astrocytes (cytoplasm marked pink, and astrocyte nuclei Ast-yellow). (B) Microglia adhering to the PI surface interposed between the implant and non-myelinated neurites containing microtubules and astrocyte branches that invaded the platform pores. No myelinated axons were seen in the immediate vicinity of the implant at this point in time after PPMP implantation. PI-polyimide ribs, gMμE-yellow asterisks, mAx-myelinated axons (green), CL -conducting line, μG-microglia, μT-microtubule, Ast-astrocyte. Note that an unmarked copy of this figure is presented as Supplementary Figure S3.

**Figure 4.**
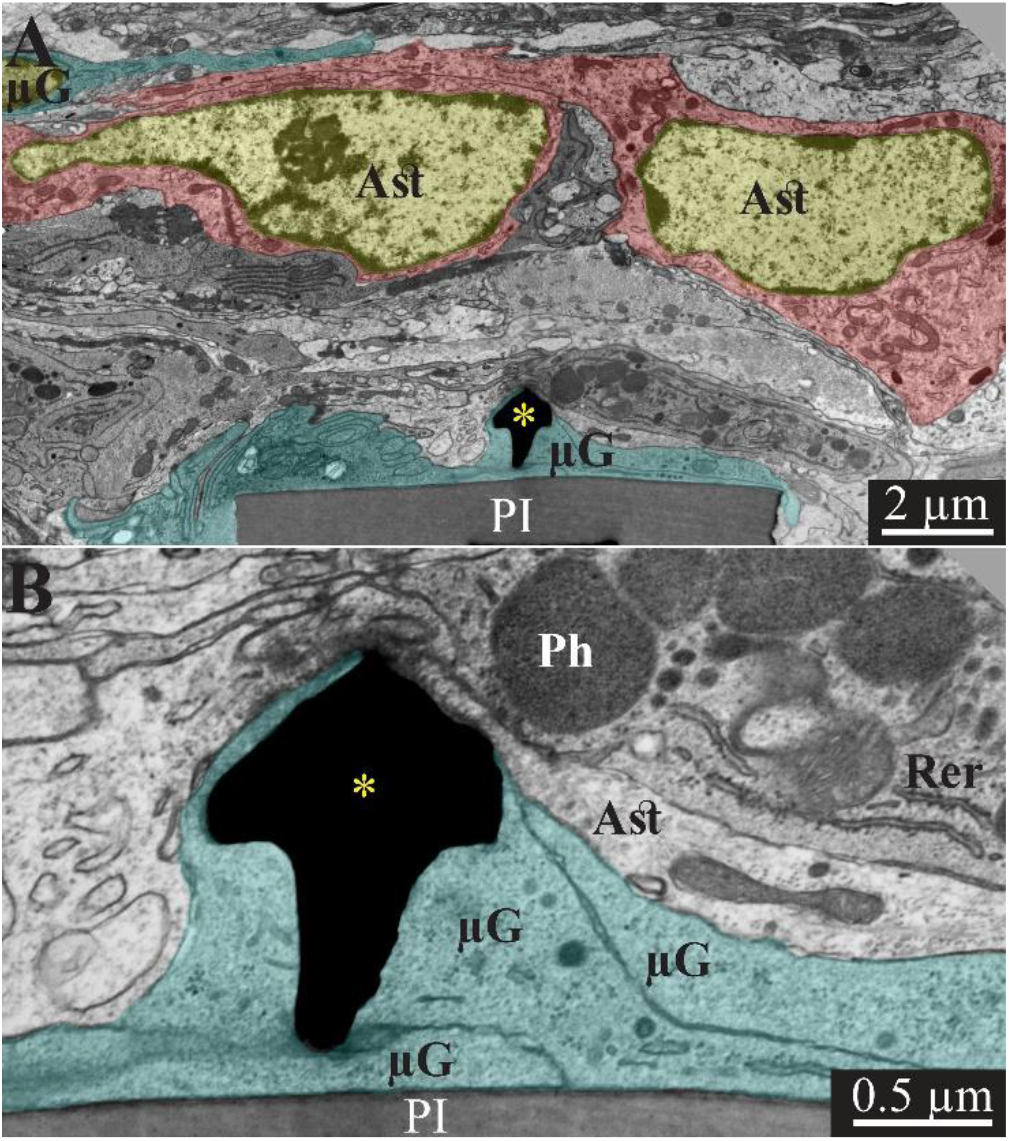
Transmission electron microscope images of a gMμE (black mushroom shaped profiles A, B) tightly engulfed by microglia, 2 weeks after PPMP implantation. Note the thin 0.5-1 μm layers of microglia branches that adhered tightly to the PI surface and the gold mushroom microelectrode (cyan). Additional microglia layers characterized by dark cytoplasm (not labeled in color) contained a rough endoplasmic reticulum and dark inclusions which plausibly were lysosomes and lipofuscine granules. Astrocyte branches characterized by sparse electron dense material containing intermediate filaments invaded in between the microglia branches but did not form direct contact with the implant. Note that the TEM section is slightly tilted in respect to the long axis of the gMμE. For that reason the mushroom’s stalk appears to taper toward the polyimide platform and looks like it end in the tissue rather than attached to polyimide. Astrocyte cell bodies resided micrometers away from the implant. PI-polyimide ribs, gMμE-yellow asterisks, μG-microglia, Rer-rough endoplasmic reticulum, Ast-astrocyte. An unmarked copy of this figure is presented as Supplementary Figure S4.

In line with Figure 2 and our earlier immunohistological studies (Huang et al 2020; Sharon et al 2021), two weeks post-PPMP implantation, TEM images revealed the presence of astrocyte cell bodies and branches in close proximity to the platform surface and within the platform pores (Figures 3B and 4A). Astrocyte cell bodies could be identified by their pale nuclei that had a thin rim of heterochromatin and pale cytoplasm (Figures 3 and 4A). Typically, the cytoplasm of astrocyte branches are characterized by sparse electron-dense material containing intermediate filaments (glial fibrillary acidic protein, GFAP, Garcia-Cabezas et al., 2016;Nahirney and Tremblay, 2021).

Two weeks after implantation, the dark microglia cytoplasm that adheres tightly to the gMμE-PPMPs surfaces are often interposed between the PPMP surfaces and the astrocytic branches, thus mechanically preventing direct contact between the neurite and astrocytic branches and the platform surfaces (Figure 3B).

Immunohistological imaging of the neurite revealed that at two weeks post-implantation, neurites extended into the PI platform pores (Figures 2C and E3). Based on the presence of microtubules in axons and GFAP in astrocytes (Figure 3B) it was possible to differentiate between the branches of the astrocytes and the unmyelinated neurites (axons and dendrites). At two weeks post-PPMP implantation, no myelinated axonal profiles were observed in the immediate vicinity (<10 μm) to the gMμE-PPMPs. Further away from the implants (>10 μm) myelinated axons were observed (Figure 3).

Neuronal cell bodies characterized by typical round euchromatic nuclei, the presence of electron dense nucleoli and nuclear membrane invaginations were observed as close as ∼20 μm from the PI platform and onwards (see also Figure 2 E4). Chemical presynaptic terminals identified by the presence of profiles containing clusters of synaptic vesicles or chemical synapses identified by presynaptic fibers in association with typical post-synaptic densities were imaged at a distances of approximately 10 μm from the platform.

### Regenerative processes in contact and around PPMP implants at four weeks post-PPMP implantation

Immunohistological examination of the changes in cell composition and distribution within and around the implanted PPMPs 4 weeks after PPMP implantation (Figure 2) suggested that the parenchyma around the implant had undergone regenerative processes. These included: (a) a reduction in the average microglia density in the first shells around the implant, but not within the PPMP pores (Figure 2 E1 and Huang et al., 2020); (b) a significant increase in the average density of the neuronal cell bodies in the first shell around the implant, preceded by the extension of neurites towards the implant and into the PPMP pores (Figure 2 E3&E4 and Huang et al., 2020). In contrast to these regenerative processes, the astrocyte branches and cell bodies continued to increase during the fourth week post-implantation both within the PPMPs and in the first shell around it (Figure 2 E2).

The overall regenerative processes observed at the confocal microscope resolution were reflected and more finely delineated at the ultrastructural levels. TEM images revealed myelinated axons extending towards and in the vicinity of the implant surface (Figure 5). A considerable increase in the density of structurally mature chemical synapses was seen. At 4 weeks post-implantation, the PPMP’s “ribs” were no longer enwrapped by dark protoplasmic protrusions emanating from microglia cell bodies. Rather, relatively thin layers of electron opaque cytoplasm adhered to the surface of the PPMPs and the gMμEs (Figure 5). Relevant to the electrophysiological recording functions of the implant (see discussion), it is noted that the extracellular cleft formed between the microglia membranes that enwrapped the gMμE remained in the range of 10-20 nm (Figure 5B). The microglia clearly interposed between the myelinated axons extending in the electrodes’ vicinity and between pale cytoplasmic profiles of astrocytic branches and unmyelinated axons. Astrocyte branches and cell bodies, microglia and unmyelinated axonal profiles were seen occupy the pores between the PI “ribs” and directly adhere to the PI surfaces.

**Figure 5.**
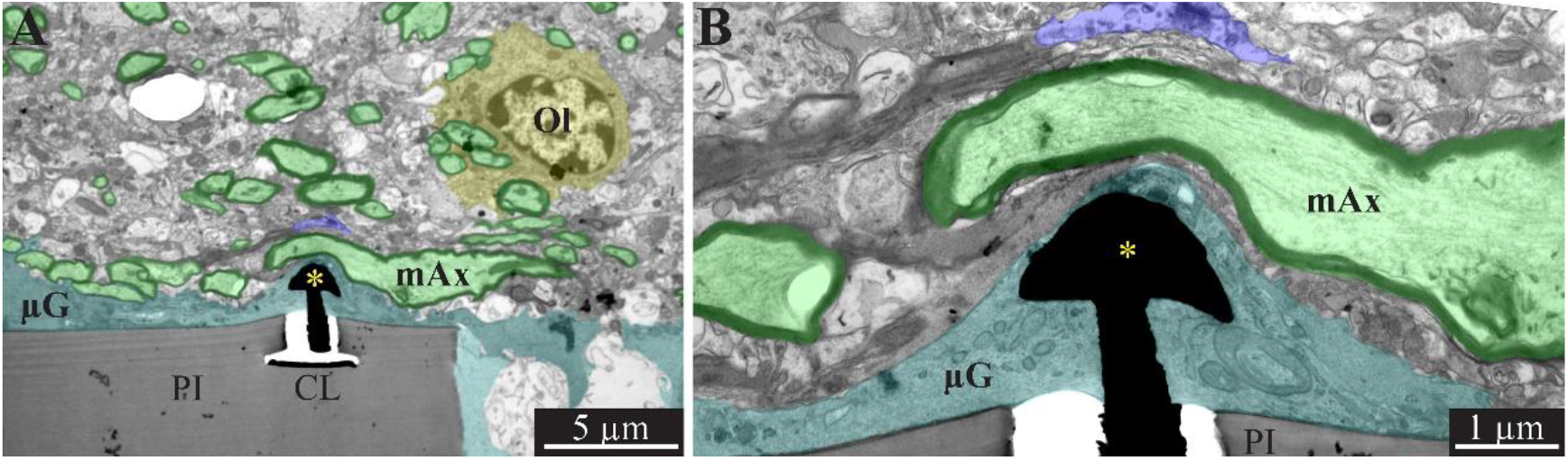
A low and high magnification, transmission electron microscope image of the interfaces formed between a gold mushroom shaped microelectrode extending from a polyimide platform implanted for 4 weeks and the surrounding cortical tissue. The mushroom shaped microelectrode (black) immerges from the PI substrate (gray). Note the thin layer of dark microglia (cyan in A and B, and unmarked dark gray branches in B) adheres tightly to the gMμE and PI substrate. The parenchyma around the implant underwent regenerative processes as indicated by the large number of myelinated axons (green and black envelope) and the presence of oligodendrocytes in the immediate vicinity of the implant (yellow). Whereas axonal branches with a relatively large diameter (∼3 μm) extended close to the gMμE, it is conceivable that the adhering microglia (and in this instance the myelin as well) insulated the electrode from the surrounding excitable tissue. PI- polyimide ribs, CL- conducting line, gMμE- yellow asterisks, Ol- oligodendrocyte, μG- microglia, mAx-myelinated axons. Note an unmarked copy of this figure is presented as Supplementary Figure S6.

In addition, large profiles of dark cytoplasm containing phagocytosed materials were occasionally observed to reside within the pores (Supplementary Figure S5). In a few cases, the section went through the nucleus of these large cells. Based on the heterochromatin distribution of the nucleus, these cells were likely to be microglia. These cell types were never observed outside the PI implant pores.

In summary, whereas clear regeneration of the neuron cell body densities, axons, dendrites and synapses took place within the first shell around the implant, dark microglia branches adhering to the gMμEs and PPMPs were still present. These adhering microglia can be assumed to electrically insulate the electrodes from the surrounding neurons (see discussion).

### Increased density of neurons near the implant surface eight weeks post-PPMP implantation

Confocal microscope imaging of the cortical parenchyma interfaced 8 weeks after PPMP implantation revealed that the overall regenerative processes that were observed 2 to 4 weeks after implantation persisted. (a) The microglia density within the implant was further reduced to half of its peak value and to a third in the first shell around the implant (Figure 2 E1 and Huang et al 2020). (b) Whereas the neurite density (NFI values, Figure 2 E3) did not change, the average neuronal cell body density in the first shell around the implant further increased to 86% with respect to the control level (Figure 2 E4). (c) On the other hand, the astrocyte (branches and cell bodies together) continued to increase mainly in the first shell around the implant (Figure 2 E2). These regenerative trends were reflected at the TEM level, in particular in that neuronal cell bodies were imaged to reside as close as ∼2 μm from the gMμE caps (Figures 6 and 7A). The narrow space between the cell bodies membrane and the gMμE caps were occupied by small profiles (with a diameter in the range of <1 μm) of either astrocytes or neurites. Parts of the gMμE stalk and caps were enwrapped by narrow (∼100 nm) dark cytoplasmic protrusions probably corresponding to microglia branches, while other parts appeared to be free of microglia. Myelinated axonal profiles were observed to form a direct contact with gMμE caps (Figure 7B). Chemical synaptic profiles were observed as close as ∼0.5 μm to the PI platform and the gMμEs, and within the parenchyma surrounding the implant. The pores within the PI platform were mainly occupied by astrocytic branches and unmyelinated axonal profiles. No synaptic structures were observed within the pores.

**Figure 6.**
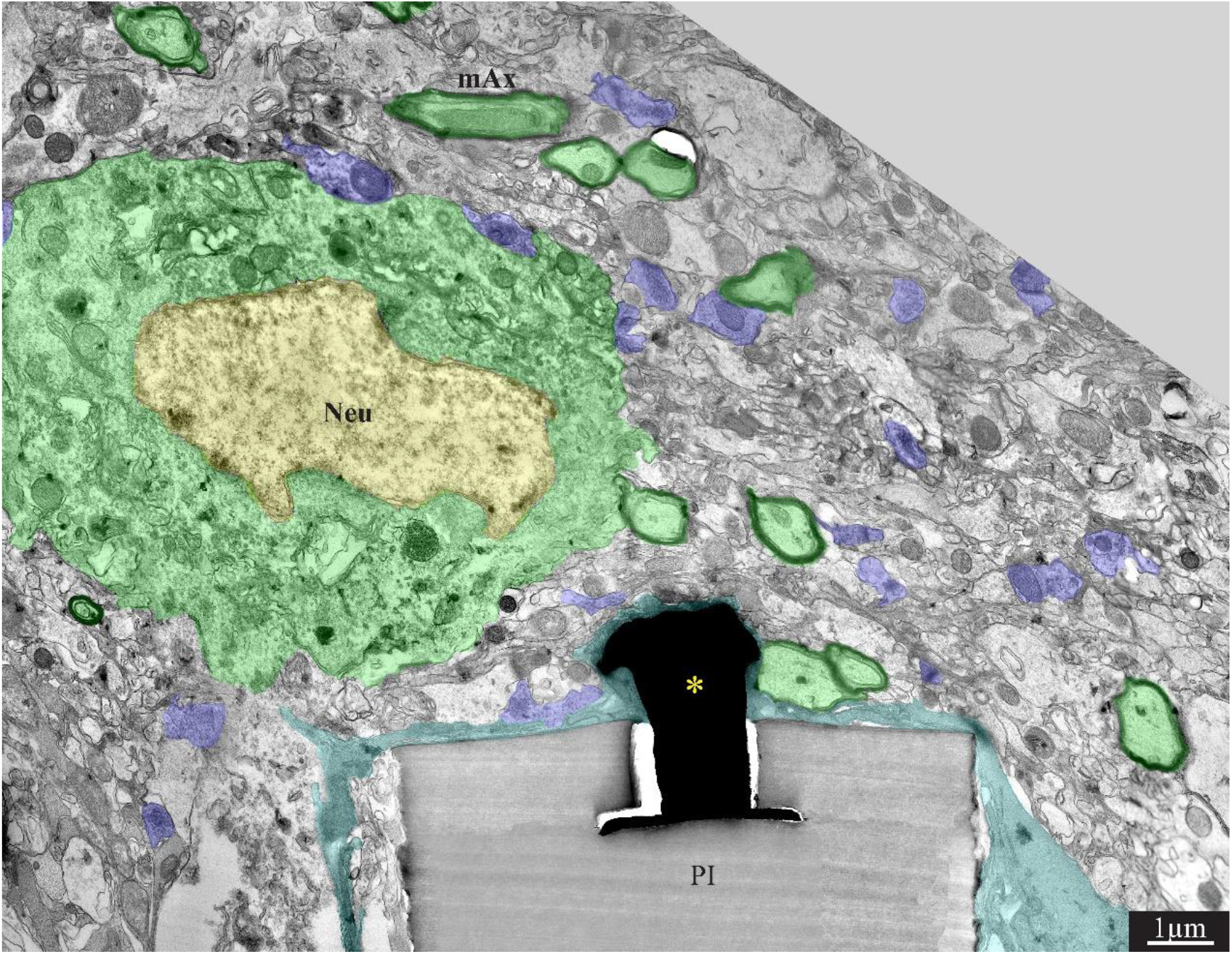
A low magnification, transmission electron microscope image of the interfaces formed between a gold mushroom shaped microelectrode extending from a polyimide platform implanted for 8 weeks and the surrounding cortical tissue. The mushroom shaped microelectrode (black) immerges from the PI substrate (gray). A thin layer of dark microglia (cyan) adheres tightly to the gMμE and the PI substrate. A neuronal cell body with a typical nuclear structure (yellow) and cytoplasm (green) resides approximately a micrometer away from the gMμE and the PI platform’s surface. Myelinated axons (green surrounded by a black sheath) are distributed in the parenchyma in contact with the microglia that adheres to the platform. Unmyelinated neurites and synaptic structures (labeled purple) were identified (using large magnification of the image) by the presence of presynaptic vesicles. The remainder of the unmarked profiles are astrocyte branches and non-myelinated neurites. PI-polyimide ribs, gMμE-yellow asterisks, Neu-neuron, mAx-myelinated axons. Note that an unmarked copy of this figure is presented as Supplementary Figure S7.

**Figure 7.**
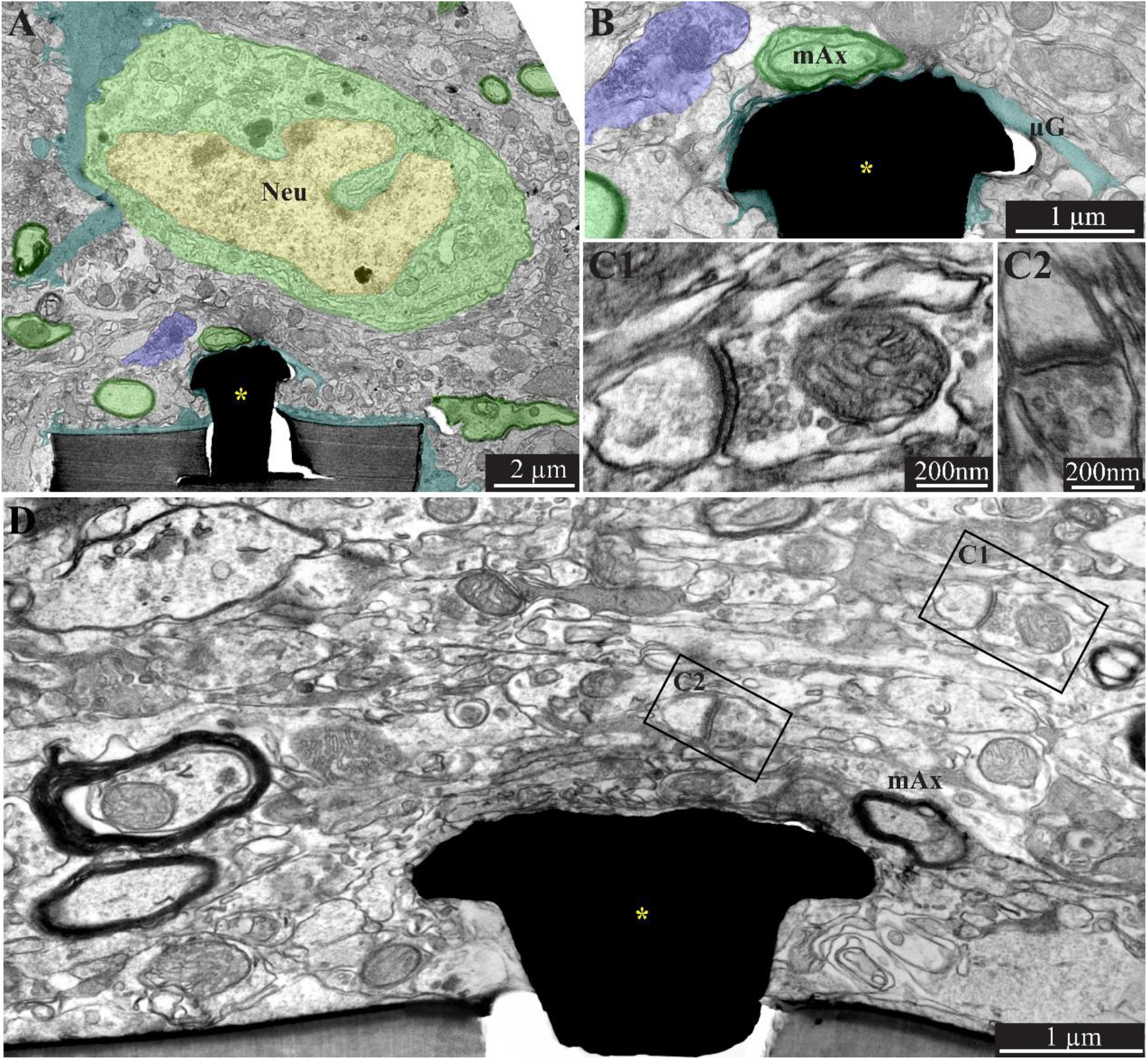
A low (A) and high magnification (B), transmission electron microscope image of the interfaces formed between a gold mushroom shaped microelectrode extending from a polyimide platform implanted for 8 weeks and the surrounding cortical tissue. As the regenerative processes of the brain parenchyma proceed with time, the dark microglia adhering layer becomes thinner (A and B, cyan). It is conceivable that even a thin microglia layer might insulate the electrodes from the surrounding parenchyma. (D) The regenerative processes of the parenchyma are also evidenced by the presence of a chemical synaptic profile as close as a few micrometers from the implant (C1 and C2, see D for the location of the synapses with respect to the electrode). Note that the TEM section (in D) is slightly tilted in respect to the long axis of the gMμE. For that reason the base of the mushrooms stalk appears to taper towards the polyimide platform. Interestingly, eight weeks after implantation we also observed gMμE that were not enwrapped by microglia and formed a direct contact with the small profile of astrocyte and possibly neurons (D). gMμE-yellow asterisks, Neu-neuron, mAx-myelinated axons, μG-microglia. An unmarked copy of this figure is presented as Supplementary Figure S8.

TEM observations conducted 8 weeks after the PPMP implantation, occasionally revealed gMμE that were not insulated by microglia. Under these conditions, the gMμEs formed a direct contact with many small (∼100 nm) axonal or astrocytic profiles (Figure 7 D). It is conceivable that the small surface area of these tentatively identified unmyelinated axonal profiles were too small to generate sufficient current to be measured by the gMμE system.

## Discussion

Despite significant progress, contemporary in-vivo MEA technologies suffer from inherent limitations that include a low signal-to-noise ratio, low source resolution and deterioration of the recording yield and FP amplitudes within days to weeks of implantation. Whereas these drawbacks constitute a critical impediment to the progress of basic and clinically oriented brain research, the mechanisms that generate these limitations remain elusive. For that reason, attempts to develop effective methods to overcome these drawbacks have only been marginally successful.

To achieve a better understanding of the mechanisms that limit the functions of implanted electrophysiological neuroprobes, for the first time, the present study examined the intact ultrastructural interfaces formed between the cortical parenchyma and a large footprint implanted neuroprobes. The findings reveal remarkable tissue regeneration around and in contact with the large-footprint implanted MEA platform. This include the regrowth of neurites towards the implant, myelination of the newly grown axons, the formation of structurally mature chemical synapses, the recovery of neuronal cell body densities in the vicinity of the electrodes (at a distance of ∼1 μm from the electrodes’ surfaces) and cortical capillaries (Figure 8). Along with this remarkable tissue regeneration, we documented that individual microglia adhering to the gMμEs-PPMP surfaces formed a micrometer-thin barrier in contact with the PI backbone and the gMμEs which we dub the “microglia-insulating-junction”. For a period of approximately 8 weeks post-PPMP implantation (the longest observation period made here), the adhering microglia prevented the formation of a direct contact between the axons or neuronal cell bodies and the gMμE. Thus, engulfment of gMμEs and most likely other 3D or planar microelectrodes by neurons is likely to be impeded. Because the microglia insulating junctions are formed at the electrode surfaces, this configuration offers an explanation to the enigma as to why no correlation has been found between the dimensions and density of the FBR and recording qualities (Kozai et al., 2014;Kozai et al., 2015;McCreery et al., 2016;Du et al., 2017;Salatino et al., 2017a;Michelson et al., 2018).

**Figure 8.**
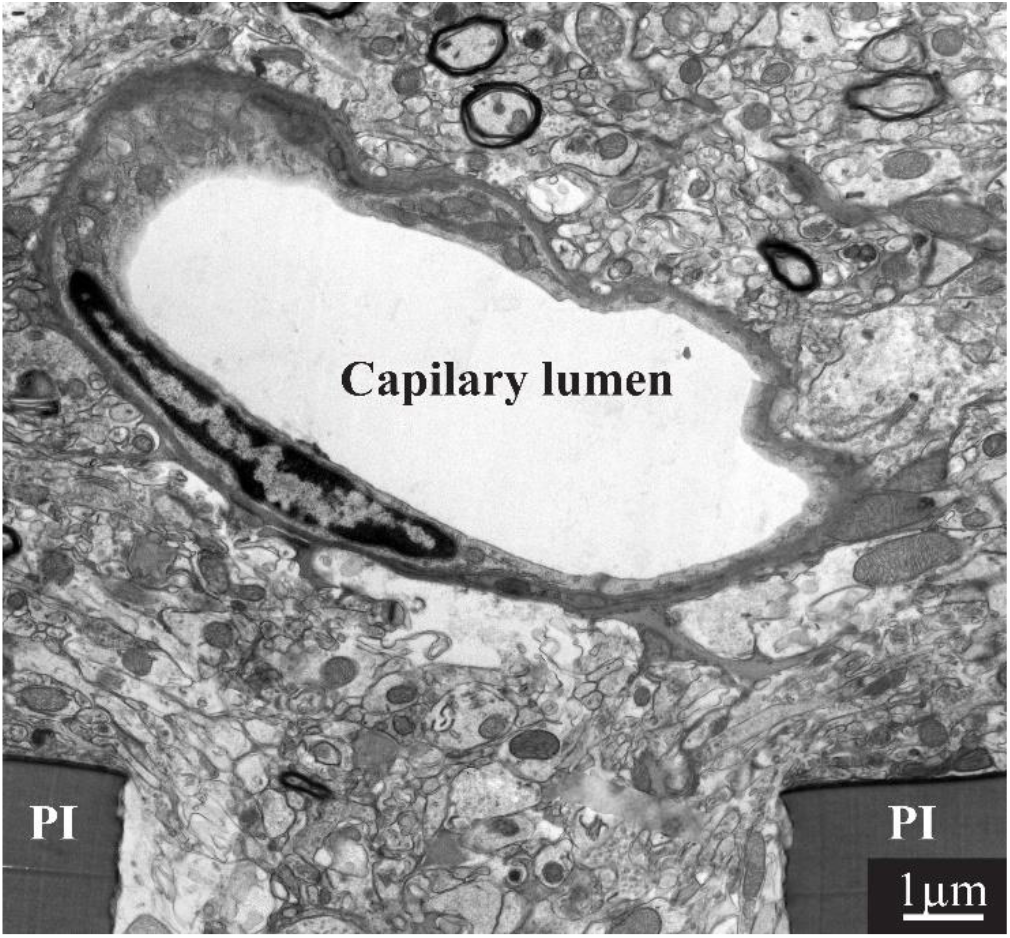
Regeneration of capillaries close to implanted PPMPs. A capillary located micrometers away from the surface of the polyimide (PI) “ribs” of an implanted PPMP for 8 weeks.

Ultrastructural analysis of the junctions formed between different cell types and planar or 3D microelectrodes under in-vitro conditions have served a pivotal role in deciphering the biophysics and potential applications of the junctions formed. An order of magnitude estimate of microglia-gMμE-junction impedance can be derived using a passive electrical circuit model composed of two parallel resistors: the seal resistance (R_s_) formed by the cleft between the plasma membrane of the microglia and the surface of the gMμEs, and the input resistance (R_μg_) of the adhering microglia.

The seal resistance (R_s_) is given by R_seal_=ρ_s_·δ /d, where ρ_s_ is the resistivity of the electrolyte solution (ρ_s_ = 0.7 ΩCm), d is the average cleft width between the neuron’s plasma membrane and the electrodes’ surface, and δ is the overlapping surface coefficient that takes into account the percentage of the electrodes’ sensitive area in contact with the microglia (Massobrio et al., 2016). Because of the unavoidable ∼20% shrinking artifact of the extracellular spaces due to the chemical fixation of the tissue for TEM imaging (Korogod et al., 2015;Hrabetova et al., 2018;Soria et al., 2020) the actual width (d) of the clefts formed between the microglia and the implanted gMμEs-PPMPs surfaces cannot be extracted with precision from the ultrastructural images. In addition, the fraction of the surface area of the contact between a gMμE or planar electrode and the adhering microglia (δ) cannot be obtained from classical TEM images. Nonetheless, an order of magnitude estimate of R_s_ formed by different cell types can be obtained by using parameters published previously in the literature. A large number of in-vitro studies have revealed that the cleft width formed between different cultured cell types and artificial substrates ranges from 20 to 100 nm (Braun and Fromherz, 1998;Iwanaga et al., 2001;Straub et al., 2001;Lambacher and Fromherz, 2002;Brittinger and Fromherz, 2005;Gleixner and Fromherz, 2006;Wrobel et al., 2008) and the contact surface area of these junctions has been estimated. The estimated seal resistance derived in these studies ranged from ∼1 MΩ in the case of planar electrodes (Weis and Fromherz, 1997;Buitenweg et al., 1998;Buitenweg et al., 2002) to ∼ 40-100 MΩs for gMμEs (Hai et al., 2009a;Fendyur et al., 2011;Spira and Hai, 2013;Ojovan et al., 2015;Shmoel et al., 2016;Massobrio et al., 2018;Spira et al., 2019).

The input resistance of mice microglia (R_μg_) was reported to be 2-5 GΩ (Avignone et al., 2008;Schilling and Eder, 2015). Since the morphology and physiology of microglia are known to change under different functional states and in response to different substrates (Eder, 1998;2005;2010;Kettenmann et al., 2011), it is conceivable that the input resistance of microglia adhering to the gMμEs is less than 2-5 GΩ. Assuming that the input resistance of adhering microglia is reduced to the range of 10-100 MΩ, the resistance formed by an adhering “microglia insulating junction” is in the range of ∼1MΩ for a planar electrode and ∼50 MΩs for a gMμE or a vertical nano-pillar engulfed by a microglia (R_μg_·R_s_/R_μg_+R_s._).

Given that the estimated resistance of intact brain parenchyma is in the range of 1-4 Ω (Logothetis et al., 2007), and 300-6000 Ω, across an encapsulation glial scar (Turner et al., 1999;Szarowski et al., 2003;Moffitt and McIntyre, 2005;Grill and Mortimer, 2014), the current generated by neurons positioned very close or in contact with microglia adhering to a sensing electrode is expected to be attenuated by 1-3 orders of magnitude. Thus, the FPs generated by neurons positioned in the immediate vicinity of a microglia-insulating-junction might be below the level of detection.

It is worth noting that recent progress in bioengineering has led to the implementation of ultra-small and ultra-flexible platforms, with dimensions comparable to those of a single neuron (Xiang et al., 2014;Fu et al., 2016;Luan et al., 2017;Zhao et al., 2017;Wei et al., 2018;Guan et al., 2019;Yang et al., 2019;Zhang et al., 2021). Immunohistological observations have shown that these ultra-small, flexible implants integrate seamlessly with brain tissue, and that under these conditions neuronal cell bodies are seen to reside in close proximity to the implant (Fu et al., 2016;Luan et al., 2017;Zhou et al., 2017;Hong et al., 2018;Yang et al., 2019). Despite the fact that the impedances of these ultra-small and ultra-flexible electrodes are similar to those of conventional implants (0.5-1MΩ) and despite the seamless integration of these platforms with brain tissue, the recorded amplitudes of the FP have been within the range of those recorded by implants that trigger FBR. These observations are inconsistent with the prevailing hypothesis that in the absence of a histological FBR the FPs amplitudes should be larger. This apparent paradox may be resolved by assuming that even if a “classical” FBR is not imaged as having been formed by these implants, microglia insulating-junctions that were not detected by standard immunohistology nevertheless formed and insulated the electrodes.

Overall, the present study resolves two critical questions: (1) what are the cellular mechanisms that underlie the limited electrophysiological functions of implanted in-vivo neuroprobes and (2), can the successfully developed and advantageous gMμE or other 3D vertical nano-pillars be applied to in-vivo settings? We posited that the insulation formed by individual microglia that tightly adhere to or engulf in-vivo implanted electrodes rather than multicellular FBR deteriorate the electrical coupling coefficient between the neurons and the implanted electrodes. The microglia electrode junctions structurally isolate and electrically insulate the electrodes from the neurons and hence limit the electrophysiological functions of the electrodes. Overcoming the challenging microglia insulating-junction requires developing new protocols to specifically and temporally target the adhering microglia. This should be complemented by methods to increase the density of neuronal cell bodies to enable the formation of direct contact with the electrodes.

It is conceivable that the effective structural regeneration of the parenchyma in the immediate vicinity of the gMμE-PPMP implants and the implant-parenchyma integration documented here reflect compound abiotic and biotic factors. For that reason, it is premature to extrapolate the observation made here to implants composed of different materials, with different microarchitecture, sizes and shapes, and implanted in different brain regions in different organisms.

## Conflicts of interest

The authors declare that the research was conducted in the absence of any commercial or financial relationships that could be construed as a potential conflict of interest.

## Author Contributions

A.S. implanted the platforms and processed the tissues for both immunohistological and transmission electron microscope sectioning together with H.E.. A.S. and H.E. analyzed the images. N.S. headed the fabrication of the perforated polyimide MEA platforms. M.M. J. supervised the PPMP implantations. Y.F. thin sectioned the tissues for TEM imaging and helped with the analysis. M.E.S. conceived, designed, and supervised the project.

## Funding

This work was supported by the Israel Science Foundation grant number 1808/19. Part of this work was conducted at the Charles E. Smith and Prof. Joel Elkes Laboratory for Collaborative Research in Psychobiology. This study is based on an earlier research project supported by the National Institute of Neurological Disorders and Stroke of the National Institutes of Health under Award Number U01NS099687.

## Acknowledgment

We thank Drs. Shimon Eliav, <u>Galina Chechelinsky</u>, <u>Maurice Saidian</u>, <u>Evgenia Blayvas</u> from the Harvey M. Kruger Family Center for Nanoscience for taking part in the fabrication of the perforated polyimide-based MEA platforms. The content is solely the responsibility of the authors and does not necessarily represent any official views of the granting agencies.

## Notes

### Competing Interest Statement

The authors have declared no competing interest.

