## Supplementary for "Ultrastructural analysis of neuroimplant-parenchyma interfaces uncover remarkable neuroregeneration along-with barriers that limit the implant electrophysiological functions"

### Supplementary Material

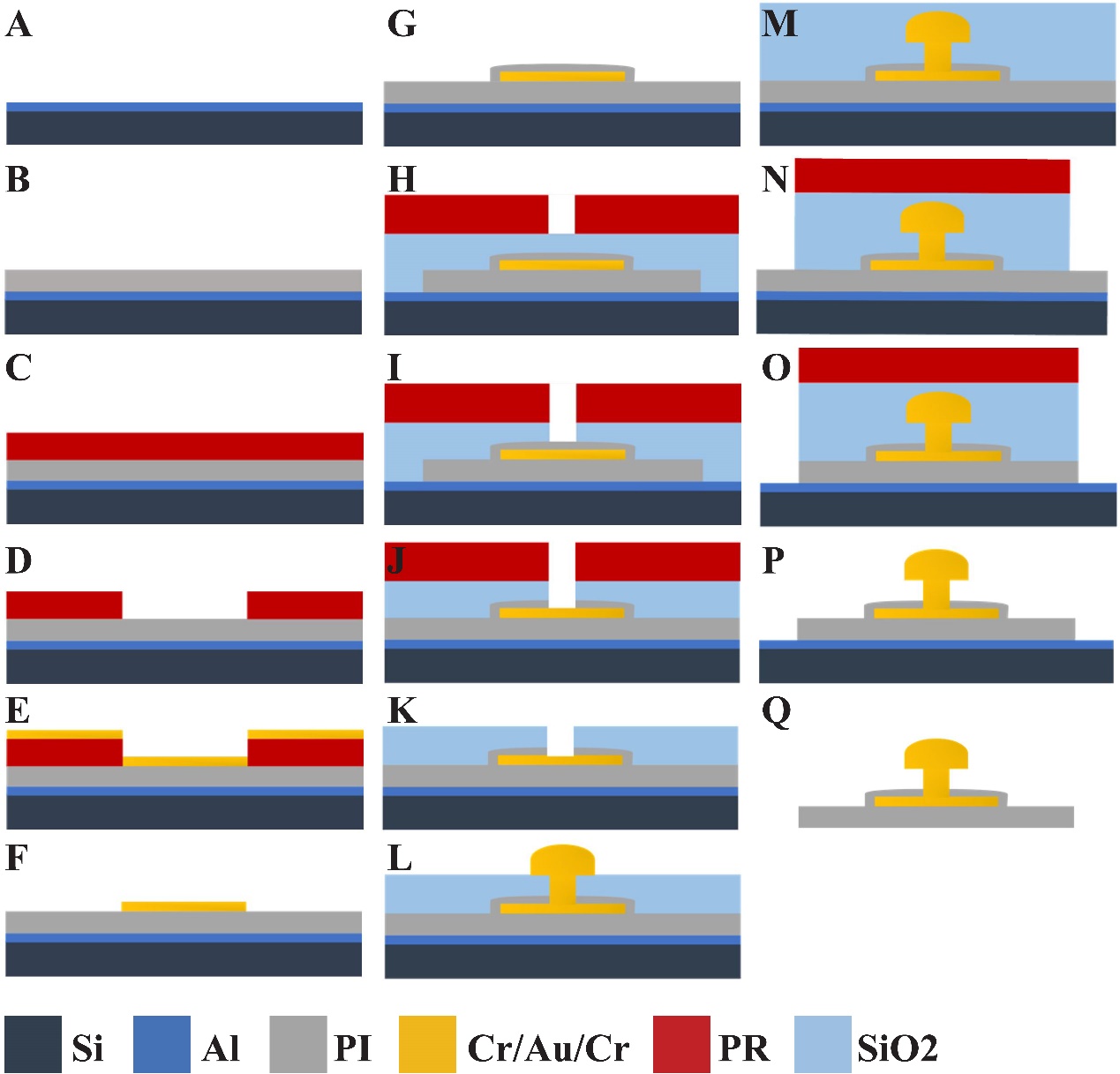
**Supplementary Figure S1. Process flow: (A)** Sputtering of a 1µm Aluminum layer as a sacrificial layer on a carrier silicon wafer. **(B)** 15µm Polyimide coating layer (3 layers of PI 2610). **(C)** and **(D)** Photoresist (nLof) coating for the 1^st^ mask to define the Cr/Au/Cr  conducting lines, pads and scribe lines. **(E)** 20nm Cr/150nm Au/20nm Cr deposition by electron beam evaporator (adhesion layer/gold conduction layer/protection layer). **(F)** Cr/Au/Cr conducting lines and pads deposition (defined by first lithography and lift off). **(G)** Spin coating of a 1000nm thick polyimide layer to encapsulate the Au conducting lines. **(H)** 20nm/1100nm Si_3_N_4_/SiO_2_ PECVD deposition,  followed by hole (via) formation and pads exposure by lithography.  **(I)** Si_3_N_4_ and SiO_2_ RIE (dry etch) (CHF_3_/SF_6_) to complete the via through the SiO_2_. **(J)** Polyimide RIE (CF_4_/O_2_) and **(K)** Cr wet etch and photoresist striping. **(L)** Electroplating. **(M)** 300nm SiO_2_ PECVD deposition. **(N)** Photolithography for the final probe etching process. **(O)** Dry etch process, Si_3_N_4_/SiO_2_ (CHF_3_/SF_6_) and Polyimide RIE (O_2_). **(P)** Photoresist striping and Si_3_N_4_/SiO_2_ wet etch (BOE). **(Q)** Remove the Al sacrificial layer to release the final devices by anodic metal dissolution.

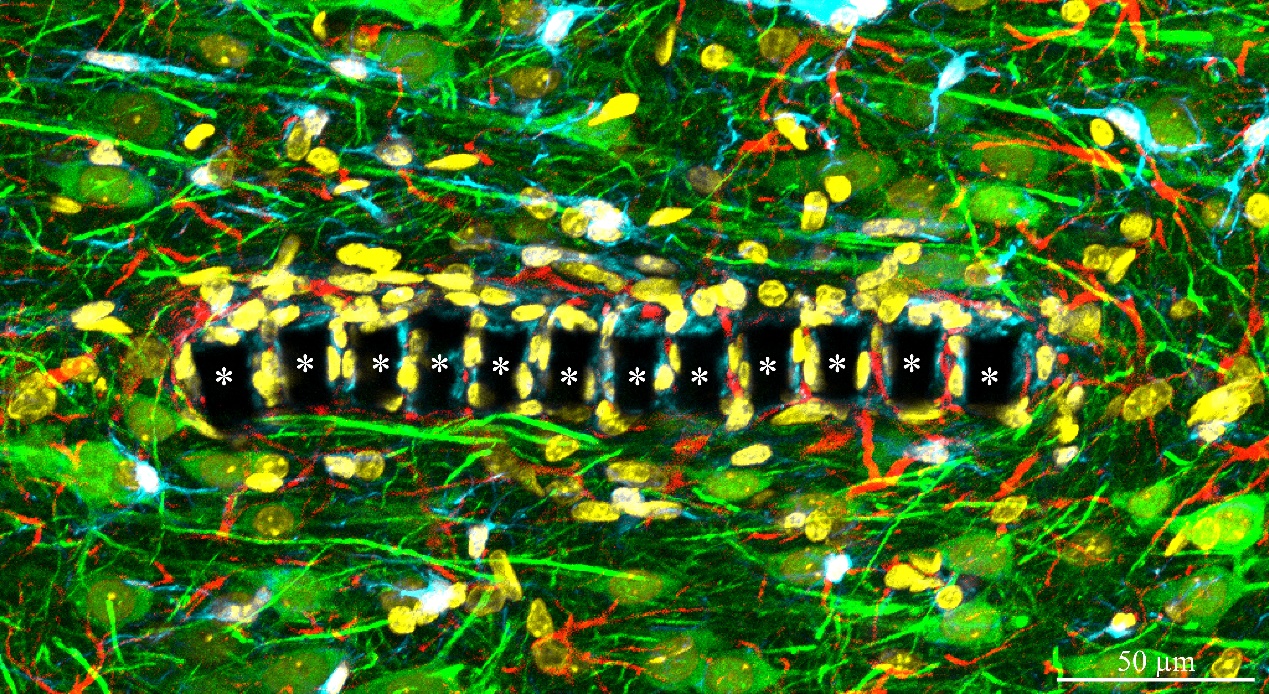
**Supplementary Figure S2.** Confocal microscope images showing a cross section through a perforated segment of an implanted PPMP along with immunolabeled cortical brain tissue, two weeks post implantation (enlargement of text figure 2D). The polyimide "ribs" (in between the pores) are indicated by asterisks. Microglia (cyan), astrocytes (red), neuronal cell bodies and neurites (green), cell nuclei (yellow).

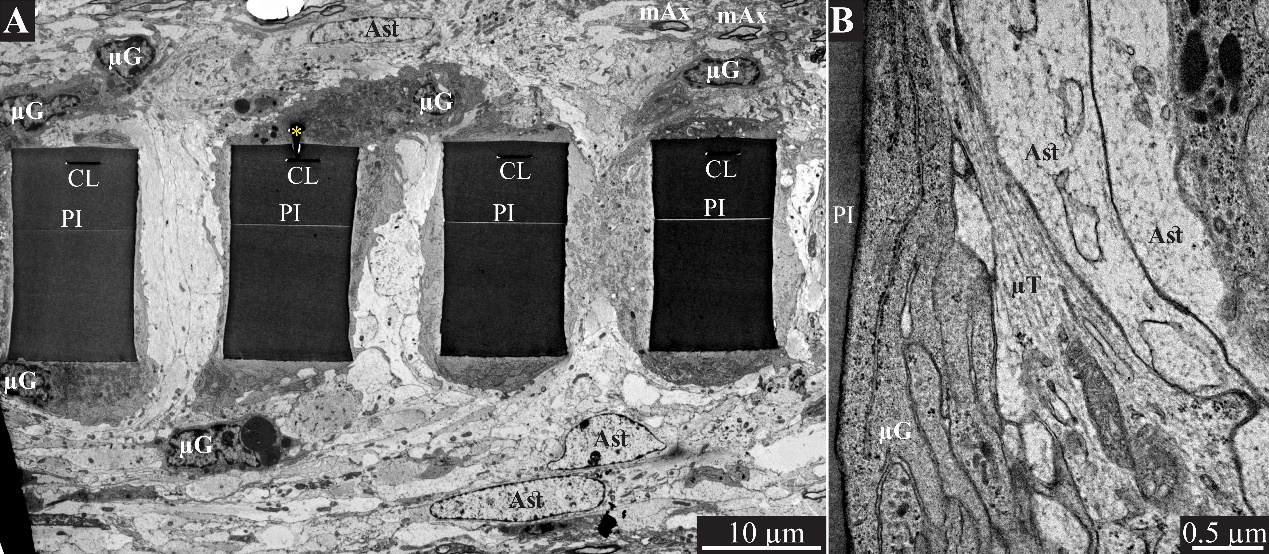
**Supplementary Figure S3. Encapsulation of induvial PPMPs “ribs” by adhering microglia.** (A) A low magnification transmission electron microscope cross-section of an implanted PPMP along with the parenchyma around it 2 weeks after the platforms implantation. The PI "ribs" are encompassed by microglia (dark cytoplasm and typical microglia nuclei- µG) and astrocytes (cytoplasm and astrocyte nuclei-Ast). (B) Microglia (µG) adhering to the PI surface interposes between the implant and non-myelinated neurites containing microtubules (µT) and astrocyte branches (Ast) that invade the platform pores. No myelinated axons were seen in close vicinity to the implant at this point in time after PPMP implantation. PI- polyimide ribs, gMµE- yellow asterisks, mAx- myelinated axons, CL -conducting line, µG- microglia, Ast- astrocyte. Note a color coded copy of this figure is presented as text Figure 3.

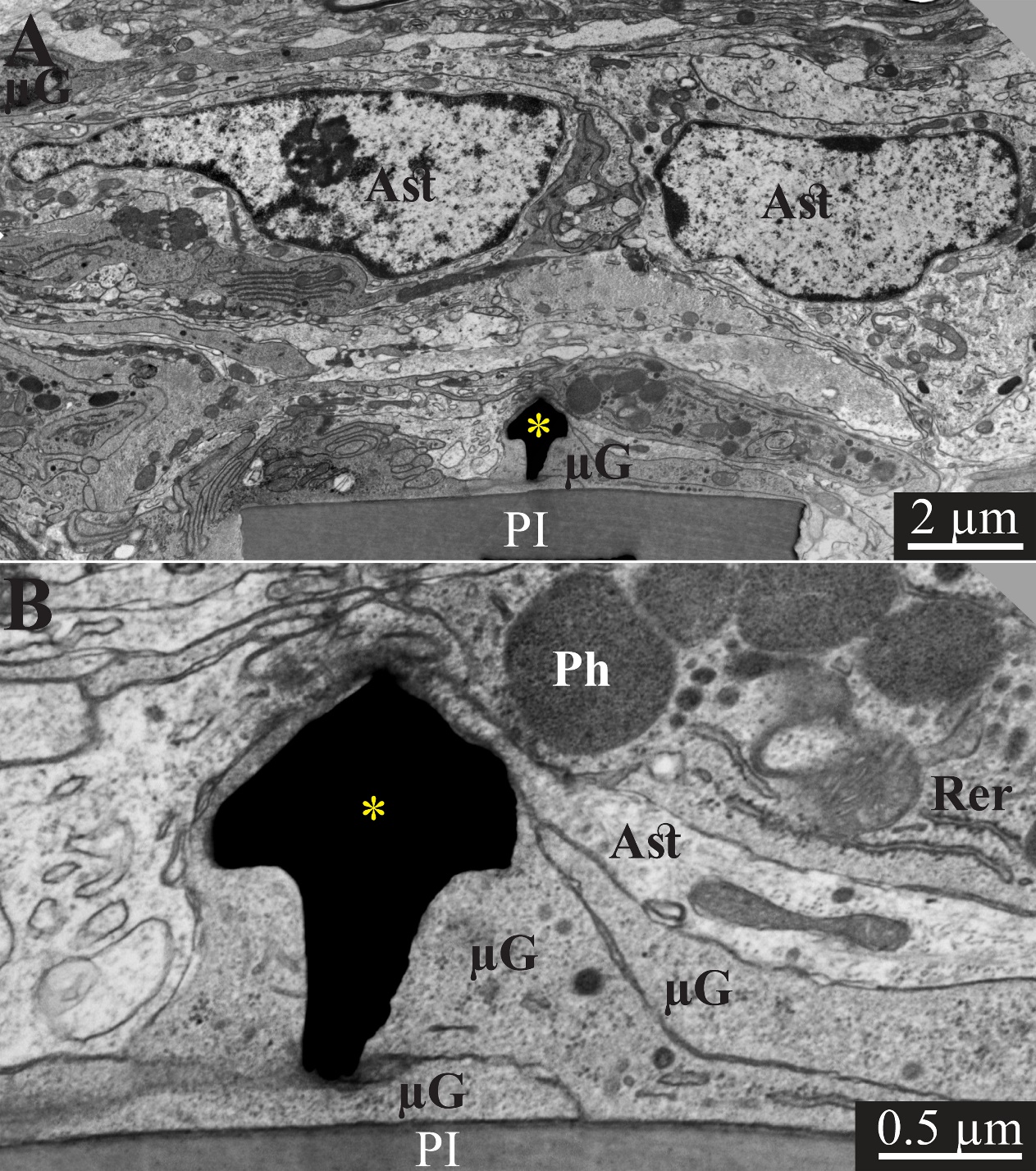

**Supplementary Figure S4.** Transmission electron microscope images of a gMµE (black mushrooms shaped profiles- asterisks) tightly engulfed by microglia, 2 weeks after PPMP implantation. Note the thin 0.5-1 µm layers of microglia branches (µG) that tightly adhere to the PI surface and the gold mushrooms microelectrode. Additional microglia layers charecterized by dark cytoplasme contain rough endoplasmic reticulom (Rer) and dark inclusions (Ph). Astrocyte branches charecterized by spars electron dense material containing intermediate ﬁlaments invade in between the microglia branches but do not form direct contact with the implant (Ast). Astrocyte cell boodies reside micrometers away from the implant. PI- polyimide ribs, gMµE- yellow asterisks, µG- microglia, Ast- astrocyte, Rer- endoplasmic reticulum. Note a color coded copy of this figure is presented as text figure 4.

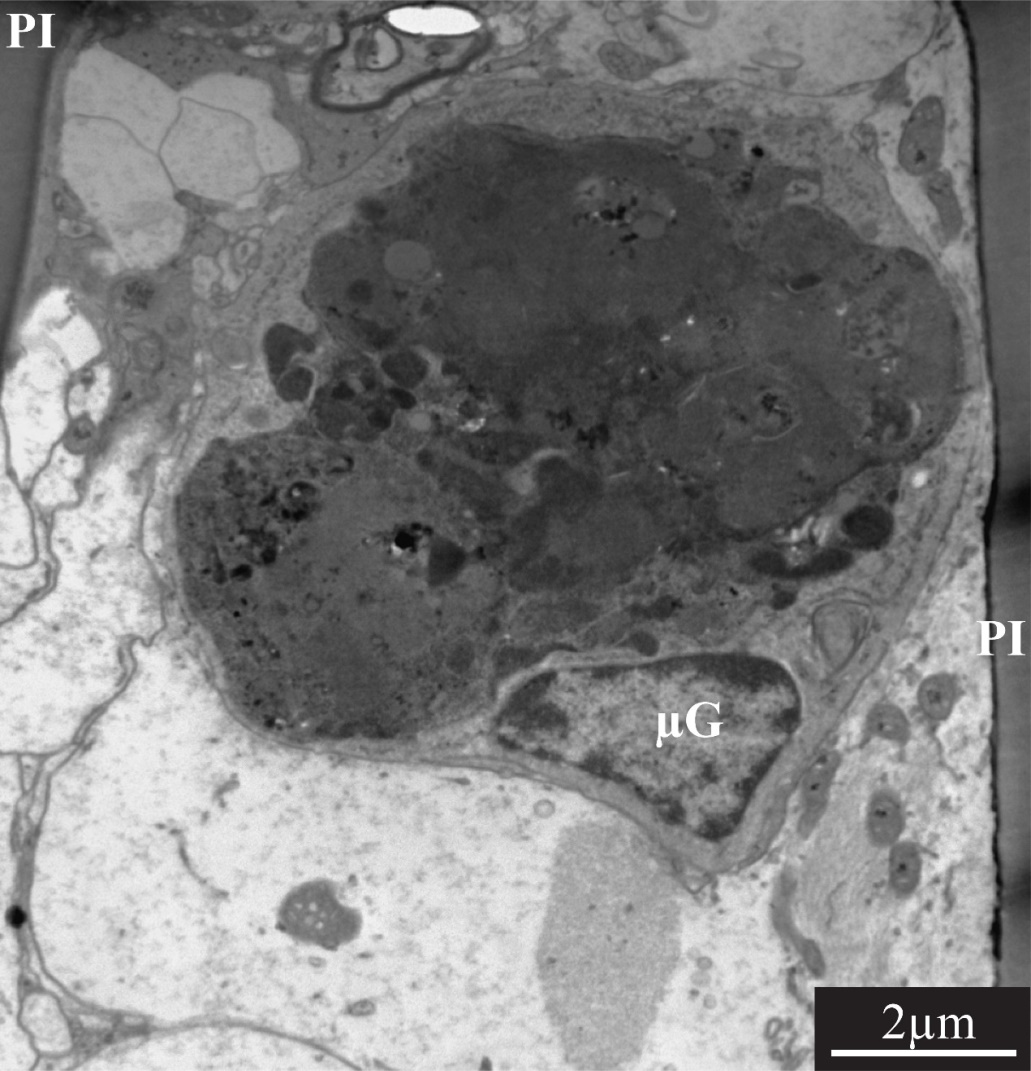

**Supplementary Figure S5.** Single large dark cytoplasm cells containing dark inclusions are occasionally seen within the perforations of PI platform implanted for 2-4 weeks. In few cases, the thin section went through the cell’s nucleus. Based on the heterochromatin distribution it is conceivable that these cells are microglia that phagocytosed cell debris at the site of implantation. These cell types were never observed outside of the PI implant pores. PI- polyimide ribs, µG-microglia.

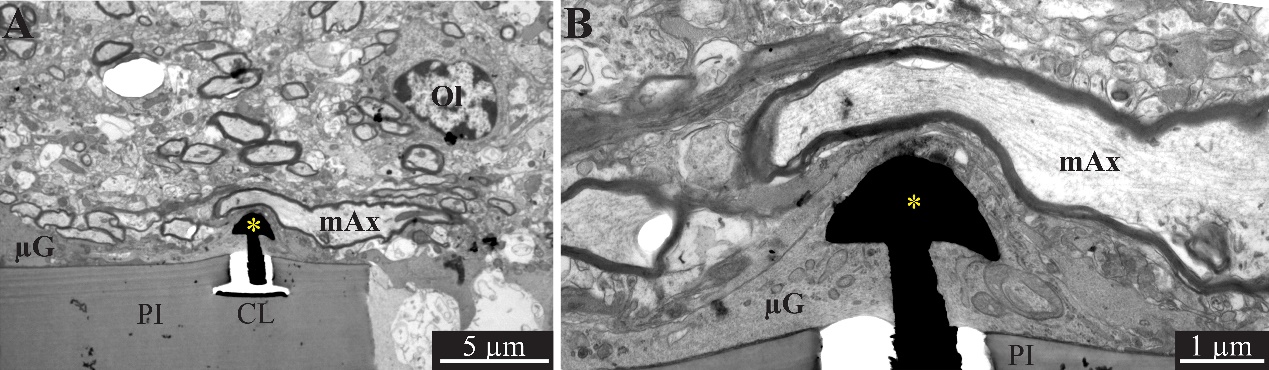
**Supplementary Figure S6.** A low and high magnification, transmission electron microscope image of the interfaces formed between a gold mushroom shaped microelectrode extending from a polyimide platform implanted for 4 weeks and the surrounding cortical tissue. The mushroom shaped microelectrode (black) immerges from a PI substrate (PI). Note the thin layer of dark microglia (µG) tightly adherers to the gMµE and the PI substrate. The parenchyma around the implant undergo regenerative processes as indicated by the large number of myelinated axons (mAx) and the presence of oligodendrocyte in close vicinity to the implant (Ol). Whereas relatively large diameter (~3 µm) axonal branches extend close to the gMµE, it is conceivable that the adhering microglia (and in this instance also the myelin) insulate the electrode from the surrounding excitable tissue. PI- polyimide ribs, CL- conducting line, gMµE- yellow asterisks, Ol- oligodendrocyte, µG- microglia, mAx- myelinated axons. Note a color coded copy of this figure is presented as text figure 5.

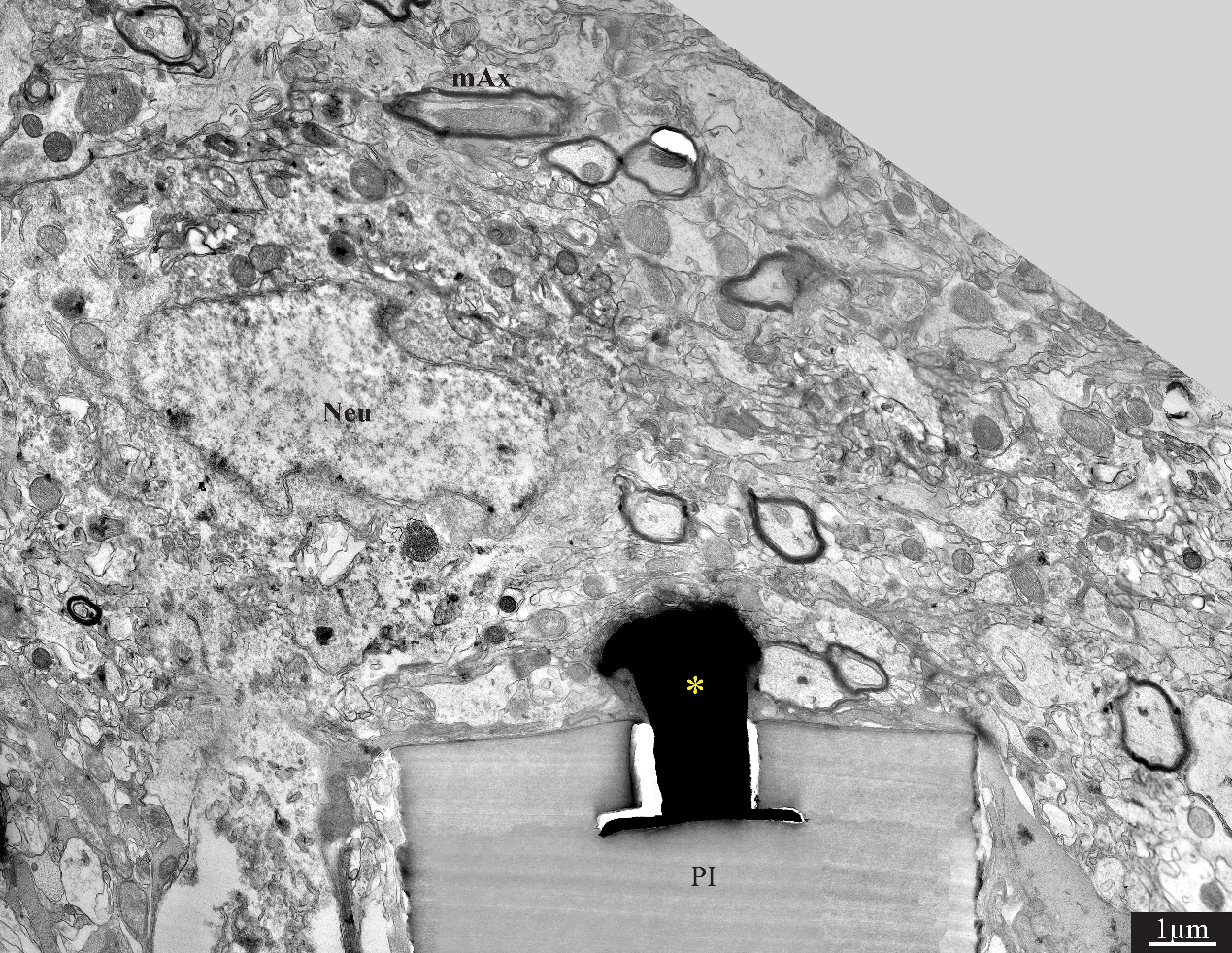
 **Supplementary Figure S7.** A low magnification, transmission electron microscope image of the interfaces formed between a gold mushroom shaped microelectrode (asterisk) extending from a polyimide platform (PI) implanted for 8 weeks and the surrounding cortical tissue. The mushroom shaped microelectrode (black) immerges from a PI substrate. A thin layer of gray microglia tightly adherers to the gMµE and the PI substrate. A neuronal cell body with a typical nuclear structure (Neu) and cytoplasm resides a micrometer away from the gMµE and the PI platforms surface. Myelinated axons (mAx) are distributed in the parenchyma in contact with the microglia that adheres to the platform. Unmyelinated neurites and synaptic structures identified (using large magnification of the image) by the presence of presynaptic vesicles. PI- polyimide ribs, gMµE- yellow asterisks, Neu- neuron, mAx- myelinated axons. Note a color coded copy of this figure is presented as text figure 6.

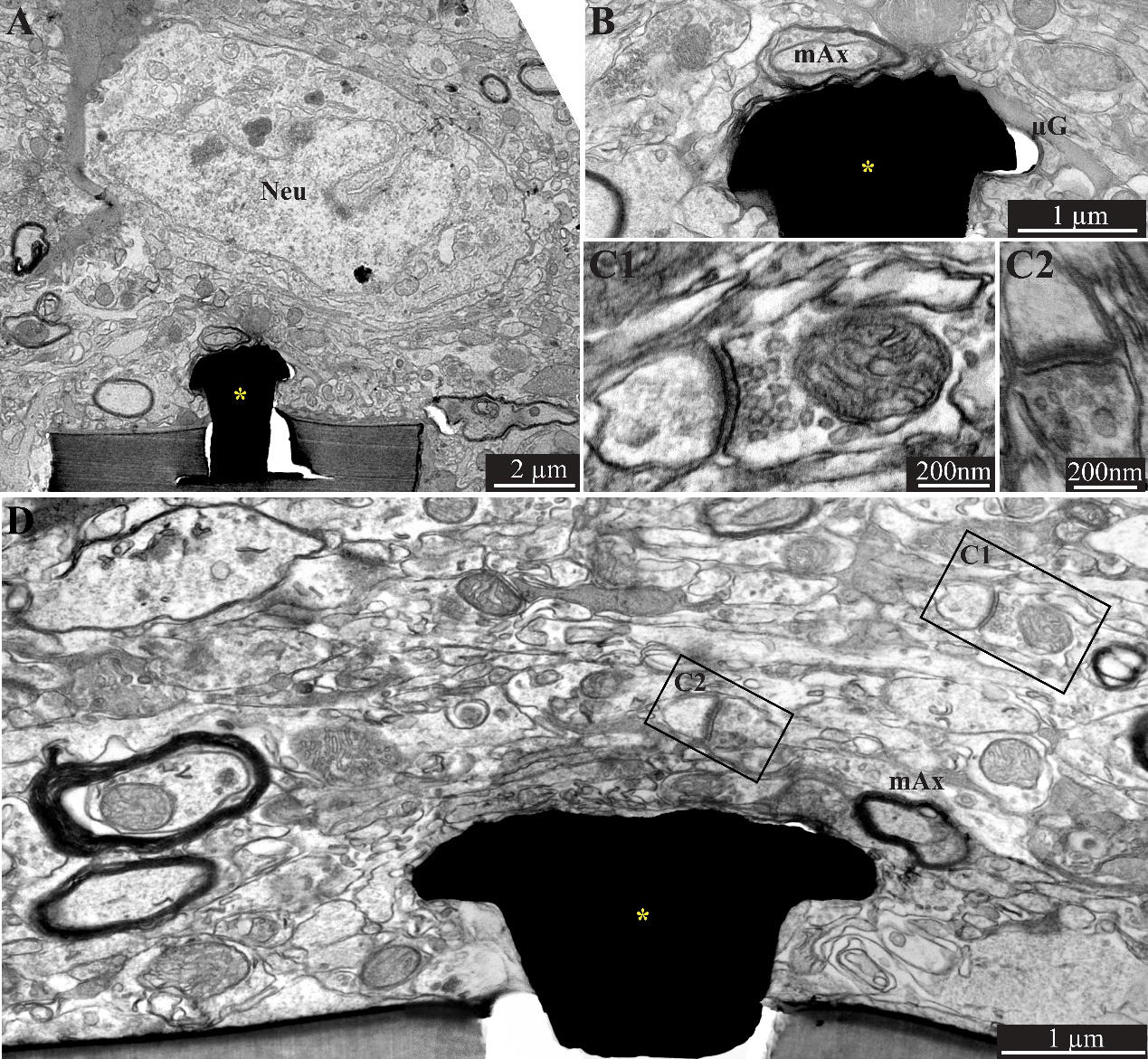
**Supplementary Figure S8.** A low (A) and high magnification (B), transmission electron microscope image of the interfaces formed between a gold mushroom shaped microelectrode (yellow asterisk) extending from a polyimide platform implanted for 8 weeks and the surrounding cortical tissue. As the regenerative processes of the brain parenchyma proceed with time, the thickness of the dark microglia (µG) adhering layer is reduced (A and B). It is conceivable that even thin microglia layer (as seen in B) might insulate the electrodes from the surrounding parenchyma. (D) The regenerative processes of the parenchyma are evidenced also by the presence of chemical synaptic profile as close as few micrometers from the implant (C1 and C2, see D for location of the synapses in respect to the electrode). Interestingly, at this point in time we occasionally observed gMµE that are not enwrapped by microglia and form direct contact with small profile of astrocyte and possibly neurons (D). Neu- neuron, mAx- myelinated axons, µG-microglia, gMµE- yellow asterisks. Note a color coded copy of this figure is presented as text figure 7.

| **Time** | **1 hour** | **1 day** | **3 days** | **1 week** | **2 weeks** | **4 weeks** | **8 weeks** |
| --- | --- | --- | --- | --- | --- | --- | --- |
| **µG*** | 17/4 | 15/4 | 13/3 | 17/6 | 15/4 | 27/5 | 19/4 |
| **As** | 19/4 | 18/4 | 10/2 | 18/7 | 29/10 | 26/5 | 18/4 |
| **Ne** | 21/4 | 18/4 | 12/3 | 16/6 | 24/7 | 22/5 | 18/4 |
| **Ne*** | 21/4 | 18/4 | 11/3 | 16/6 | 24/7 | 22/5 | 18/4 |

**Supplementary Table S1.** Number of examined brain Slices (n)/ Hemispheres (N). Ten optical sections were made per a single brain slice. µG- Microglia; As- Astrocytes and Ne- Neurons. *Cells count.

|  | 1 hour | 1 day | 3 days | 1 week | 2 weeks | 4 weeks | 8 weeks |
| --- | --- | --- | --- | --- | --- | --- | --- |
| Microglia# - Shells E | **1.97±2.67** | **2.81±1.72** | **16.01­­­±6.46**  P(1W)=0.02  P(2W)=0.03  P(4W)=0.13  P(8W)=0.012 | **20.80±6.04**  P(2W)=0.47  P(4W)=0.14  *P(8W)=2.4*10^-6^ | **20.64±6.31**  P(4W)=0.17  *P(8W)=1.8*10^-5^ | **18.57±7.25**  *P(8W)=1.8*10^-5^ | **11.19±3.2** |
| Microglia#  - Shells 1 | **2.83±0.87** | **5.37±1.06** | **16.63±3.12**  P(1W)=0.27  *P(2W)=2.7*10^-8^  *P(4W)=1.8*10^-8^  *P(8W)=1.7*10^-9^ | **17.47±4.15**  *P(2W)=9.8*10^-9^  *P(4W)=6.9*10^-9^  *P(8W)=2.2*10^-10^ | **7.84±2.93**  P(4W)=0.35  *P(8W)=9.3*10^-4^ | **8.20±3**  *P(8W)=4*10^-6^ | **4.90±1.12** |
| Astrocytes NFI- Shells E | **0.26±0.14** | **0.2±0.14** | **0.54±0.25**  *P(1W)=2.3*10^-4^  *P(2W)=5.1*10^-7^  *P(4W)=3.4*10^-9^  *P(8W)=1.4*10^-9^ | **1.04±0.41**  *P(2W)=0.007  *P(4W)=8.8*10^-7^  *P(8W)=2*10^-6^ | **1.46±0.74**  *P(4W)=2.3*10^-4^  *P(8W)=4.5*10^-3^ | **2.49±1.18**  P(8W)=0.04 | **2.01±0.61** |
| Astrocytes NFI- Shells 1 | **0.68±0.19** | **0.58±0.16** | **1.04±0.42**  P(1W)=0.014  *P(2W)=2.4*10^-3^  *P(4W)=4.6*10^-7^  *P(8W)=6.2*10^-8^ | **1.41±0.31**  P(2W)= 0.09  *P(4W)=5*10^-6^  *P(8W)=1.1*10^-6^ | **1.62±0.75**  *P(4W)=1.4*10^-3^  *P(8W)=2.7*10^-5^ | **2.26±0.77**  P(8W)=0.04 | **2.72±0.82** |
| Neurons NFI- Shells E | **0.36±0.21** | **0.16±0.14** | **0.19±0.15**  P(1W)=0.28  P(2W)=0.05  P(4W)=0.22  P(8W)=0.08 | **0.16±0.09**  *P(2W)= 0.005  P(4W)=0.02  *P(8W)=0.005 | **0.29±0.22**  P(4W)=0.1  P(8W)=0.29 | **0.23±0.11**  P(8W)=0.17 | **0.26±0.13** |
| Neurons NFI- Shells 1 | **0.74±0.15** | **0.63±0.17** | **0.67±0.15**  P(1W)=0.05  P(2W)=0.05  P(4W)=0.11  *P(8W)=1.4*10^-4^ | **0.77±0.18**  P(2W)=0.41  P(4W)=0.41  *P(8W)=0.005 | **0.79±0.28**  P(4W)=0.34  P(8W)=0.014 | **0.76±0.26**  *P(8W)=4.5*10^-3^ | **0.99±0.26** |
| Neurons# - Shells E | **0.04±0.19** | **0±0** | **0±0**  P(1W)=0.17  *P(2W)=2.3*10^-4^  P(4W)=0.03  P(8W)=0.011 | **0.04±0.16**  *P(2W)=8.8*10^-4^  P(4W)=0.05  P(8W)=0.017 | **0.41±0.5**  P(4W)=0.3  P(8W)=0.29 | **0.32±0.72**  P(8W)=0.2 | **0.55±0.93** |
| Neurons# - Shells E | **1.08 ±1.56** | **2.5±1.38** | **2.72±0.98**  *P(1W)=1.4*10^-3^  *P(2W)=1.3*10^-10^  *P(4W)=5.6*10^-9^  *P(8W)=5.5*10^-11^ | **6.33±4.01**  P(2W)=0.013  P(4W)=0.09  *P(8W)=2.5*10^-3^ | **9.13±3.12**  P(4W)=0.1  P(8W)=0.18 | **7.96±2.81**  P(8W)=0.013 | **9.94±2.55** |

**Supplementary Table S2.** Mean ± one standard derivation of the average number of cells per 100µm^2^ (# microglia and neurons) or Normalized Fluorescent Intensity (NFI) (for astrocytes and neuron cell bodies and neurites) within the implanted platform (referred to as Shell E) and 0-25µm away from the platform’s surface (Shell 1) of the perforated segment only. T-test was conducted for two-samples assuming unequal variances. P<0.01 Indicated by asterisks. 1w- 1 week, 2w- 2 weeks, 4w- 4 weeks, 8w- 8 weeks after PPMP implantation.

Bassline of microglia per 100µm^2^=2.63±0.34 (N=56).

Bassline of neurons per 100µm^2^=11.6±1.79 (N=28).
